# Horizontal transfer of an antimicrobial peptide across insects

**DOI:** 10.64898/2026.03.03.709459

**Authors:** Cedric Aumont, Pankaj Dhakad, Yi-Ming Weng, David M. Alff, Dino P. McMahon, Mark A. Hanson

## Abstract

Antimicrobial peptides (AMPs) are key defence molecules of the innate immune system. Understanding the origins of AMPs can help to explain how immune systems acquire novelty and vary in their defence capacities. However, AMPs evolve rapidly, and so the origins of similar AMPs across organisms is often unclear. Furthermore, false negatives due to low search sensitivity are common and can hinder confident annotations about true absences. As a result, understanding whether similar AMP genes found in diverse organisms represent ancestral molecules or evolutionary novelties has been challenging. Here, we present evidence of horizontal gene transfer (HGT) of the antifungal peptide gene Drosomycin across insects. We show that in Diptera, the presence of Drosomycin is restricted to the Drosophila Melanogaster species group and additionally the distant relative Drosophila busckii. We go on to recover Drosomycin genes in cockroaches (Blattodea), mantises (Mantodea), katydids (Orthoptera), beetles (Coleoptera), parasitic wasps (Hymenoptera), and a recently acquired and pseudogenized Drosomycin locus in Liposcelis booklice (Psocodea), but not other insects. Explaining this diversity through shared ancestry requires at least 60 independent loss events, or just nine HGT events. Previous studies have suggested that similar AMPs found across divergent species reflect conservation from a common ancestor, or due to their small size, that they arose via convergent evolution. Our findings suggest horizontal gene transfer can be responsible for the presence of some AMP genes scattered across the tree of life. By presenting a mechanism through which immune systems can acquire novelty, our study also suggests a possible explanation for certain lineage-specific defence competencies against infectious disease. While loss of AMP genes is well-documented, here we suggest gain of AMPs can occur just as suddenly.

## Introduction

Understanding why animals respond differently to the same infections is of major importance to the One Health approach [1]. The innate immune response is a front line defence against infection, mediated by effectors such as antimicrobial peptides (AMPs, also called “host defence peptides” or HDPs). Immune genes are among the most rapidly evolving genes in the genome across the tree of life [2–4]. In recent years, major advances have been made in our understanding of the importance of AMPs in host physiology and defence against infection [5–8]. Indeed, in mammal and insect populations, genome-wide association studies have now pinpointed variation in single AMP genes as potent determinants of susceptibility to infection by specific microbes [9,10]. This important and highly-specific activity is proposed to come from host AMP evolution targeting the “Achilles heel” of ecologically-relevant bacteria over evolutionary timescales [11], providing hosts with uniquely attuned effectors to fight specific pathogens. Understanding the origins and evolution of host AMPs is thus a promising avenue to determine some of the underlying genetic causes for individual- or species-specific susceptibility to infectious diseases.

Immune novelty is expected to arise most commonly through host immune receptor and immune effector diversification [12]. Genes can diversify for novel functions through a myriad of mechanisms including duplication (alongside subsequent neo- or subfunctionalisation), exon shuffling, retroposition, *de novo* origination, and horizontal gene transfer (HGT, also called lateral gene transfer) [13–15]. The term “antimicrobial peptide” can apply to mature proteins as short as 13aa (e.g. Indolicidin), or as long as ∼200aa (e.g. some Attacins). Because AMP genes evolve so rapidly, even seemingly-related AMPs can arise from convergent evolution of a sufficiently similar precursor molecule, exemplified by the convergent Diptericins of tephritid and drosophilid fruit flies that specifically combat *Acetobacter* bacteria present in rotting fruits [16]. The Diptericins themselves are part of the glycine-rich Attacin superfamily of AMPs [17,18]. While complex, such relationships can be inferred for these longer “peptides” of >50 residues owing to conserved motifs, even when two mature proteins are as little as ∼20% similar.

On the other hand, short AMPs often contain compositional biases (e.g. proline-rich peptides, cysteine-rich peptides). Highly similar 20-30aa proline-rich peptides exist across mammals and insect orders, including e.g. Tur1a (dolphins), Pyrrhocoricin (firebugs), Abaecin (honeybees), and Drosocin (fruit flies) [19–22]. In these cases, it is difficult to assess whether these similar AMPs harbour an ancient common ancestry, or evolved convergently towards a relatively simple sequence containing multiple prolines. Furthermore, plants and animals encode tens to hundreds of related AMPs that help to regulate the host microflora [23–27]. As a consequence of AMP duplication and subsequent diversification episodes, the chance that two AMPs from different species bear a superficial resemblance is quite high. Indeed, Defensin peptides are found in plants, animals, and fungi, typified by encoding 6 cysteine residues stapling together alpha helix and beta sheet domains. This protein structure has evolved by convergence at least once, and in the case of marine “Big Defensins”, multiple events of gene or motif loss have taken place [28–30]. It should also be noted that in certain cases, AMP genes can be widely conserved with highly similar and relatively stable mature sequence, even if individual genes are sporadically lost over evolutionary timescales [31,32]. There is thus support for both convergent evolution and ancient conservation of AMP genes. However, in some cases, strikingly similar genes can be found in distantly related taxa, where convergent evolution seems insufficient to explain their similarity, yet conservation stemming from a last common ancestor is highly non-parsimonious. Such cases have been puzzling.

Horizontal gene transfer describes the transmission of genetic material from one organism to another [33]. In insects, vectors of HGT can include parasites, bacteria, plants, fungi or viruses [14,34–38]. Horizontally transferred genes represent just a small fraction of genes in the genomes of most species [37]. While ancient HGT events will have occurred in many lineages, pinning down ancient horizontal transfer of short, rapidly evolving AMP genes can be difficult given the limitations of sequence similarity-based approaches. Moreover, potential autotoxicity of the AMP response suggests that acquiring additional foreign AMPs could even be harmful to the host [39–42], which may select against their acquisition. Whether AMPs introduced to a novel host genome would be functional in host physiology or immunity is itself unclear, as the regulatory machinery must be transferred alongside the coding sequence and must be compatible with the new host. Indeed, there are few concrete examples of horizontally transferred AMP genes outright, much less of horizontally transferred AMPs participating in the immune response of a novel host.

Here we explore the evolutionary origin of the *Drosomycin (Drs)* gene family across insects. Drosomycin was originally described in *Drosophila melanogaster* through protein isolation, and is thought to be restricted to the Melanogaster group of the subgenus Sophophora [43,44]. In *D. melanogaster,* the *Drs* gene encodes a precursor protein of 60 amino-acids, characterised by a signal peptide region and the Drs antifungal mature peptide. Drosomycins are Knottin/Defensin-like peptides but with 8 cysteine residues that staple together a cysteine-stabilized alpha-beta motif, including a bond between the 1st and 8th cysteine at the N- and C-termini of the peptide [18,45]. Surprisingly, we found *Drs* genes in a single distantly related drosophilid aly, a finding that is hard to explain by shared ancestry with the *Drs* genes of *D. melanogaster*. We further recovered highly similar *Drs* sequences from mantises, cockroaches, beetles, katydids, parasitic wasps, and even pseudogenes in booklice. However, a systematic screen of genomic and transcriptomic data from 25 winged insect orders, and outgroups, revealed no additional taxa encoding *Drs* genes. We conclude that *Drs* has undergone at least nine HGT events across the surveyed insect lineages. In at least one non-*D. melanogaster* lineage *Drs* is immune induced. Our study therefore provides a case study of multiple horizontal transfer events of an AMP gene across distantly related species. In doing so, we provide a mechanism to explain the origins of an AMP that appears sporadically across disparate taxa. This finding provides a conceptual basis for how unrelated species can rapidly acquire novel defence responses to combat infection.

## Results

### Horizontal transfer of *Drosomycin* in Drosophilidae

#### Two *Drosomycin* gene clusters are present in *Drosophila busckii*

In a previous study, Drs protein was described as present in the house aly *Musca domestica* [46]. However, within Drosophilidae, multiple studies have since confirmed *Drosomycin* and *Drosomycin-like* genes from only the Melanogaster group of the subgenus Sophophora (chromosome 3L, cytogenetic map 63D) [44,47]. In *D. melanogaster*, this gene family includes the canonical *Drosomycin* gene (*Drs*), and six Drosomycin-like genes, *Drsl1-6*. To verify the ancestral state of the *Drs* gene in Diptera, we screened the genomes of diverse aly species for the presence of *Drs* genes, which included many assemblies of chromosome-level quality [48] (Table S1).

In our screens, we found no evidence of *Drs* in any dipteran outside the family Drosophilidae; this includes *M. domestica,* for which we screened three independent genomes. Originally, *M. domestica Drs* was proposed based on a protein sequence and PCR [46], which, in light of modern genomic sequence, likely reflects contamination of that sample by *Drosophila* DNA (see supplementary text). Surprisingly, however, we did recover five *Drs-like (Drsl)* genes present in two gene clusters in the aly *Drosophila busckii* (Figure 1a). The presence of *Drs* in *D. busckii* was unexpected because *D. busckii* is allied with the subgenus Drosophila, a lineage that diverged from the Melanogaster group of the subgenus Sophophora ∼50 million years ago (mya) [49]. Yet, no subgenus Drosophila flies, nor subgenus Sophophora flies of the Willistoni, Saltans, or Obscura groups have *Drs* genes (Figure 1c). We describe these *D. busckii Drs-like* gene clusters as cluster 2L *(DbDrsl1, DbDrsl2, DbDrsl3)* and cluster 2R *(DbDrsl4, DbDrsl5).* These two *Drs* gene clusters are present in the genome assemblies of three independent *D. busckii* strains (GCA_011750605.1, GCA_001277935.1, GCA_001014355.1). The *D. melanogaster Drs* genes are located on chromosome 3L (Figure 1b). Inspection of the genomic neighbourhood (hierarchical orthologous groups “HOGs,” see [50]) of the *D. busckii Drs-like (DbDrsl)* clusters indicates that nearby genes are syntenic orthologues of genes found on *D. melanogaster* chromosome 2L (cytogenetic map 21B and 32A) and 2R (cytogenetic map 56F and 57F) (Figure 1ab, Table S2). Furthermore, these *D. busckii Drs* genes occur rather tightly between neighbouring genes, often less than 5kb apart, which suggests they were inserted through precise mechanisms (e.g. transposon insertion) and not via a larger genomic recombination event. Because the two *D. busckii* loci have genes that are more similar to their locus partner than across the genome, we consider it more likely that a single transfer of a *Drs* gene occurred first, followed by tandem duplications (ex. shown later in Figure 3). We further inspected genes from the genomic neighbourhood of *D. melanogaster Drs* on chromosome 3L (e.g. *ckd, CG12017, kst, sty*), finding most orthologues to be located on chromosome 3L of *D. busckii*. However, we did not find any *Drs-like* genes in the 3L gene regions of *D. busckii* after manually querying the translated nucleotide sequence of this 3L region for Drs protein motifs. Taken together, the *DbDrsl1-5* genes of *D. busckii* found on *D. busckii* chromosome 2 are not likely to be the result of a genomic translocation event involving an ancestral *Drs* locus common to both the Melanogaster group and *D. busckii*. Instead, genomic synteny analysis suggests the *Drosomycin-like* genes of *D. busckii* and the Melanogaster group were inserted into their genomic loci independently.

**Figure 1:**
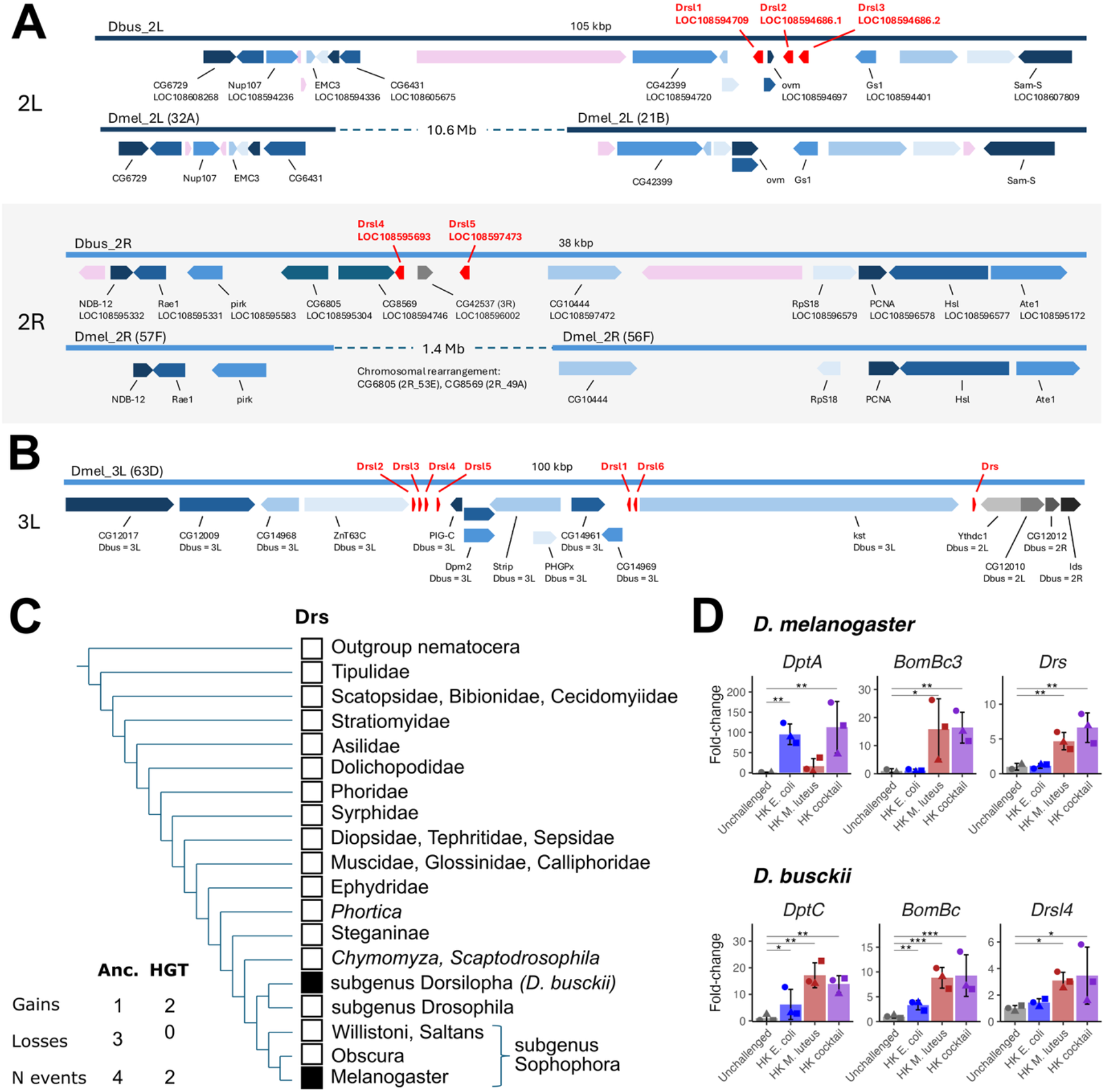
*Drosomycin* has been horizontally transferred into *Drosophila* species at least once. **A)** *Drosophila busckii* encodes five *Drsl* genes across two genomic clusters on chromosome 2 (red genes). The syntenic genome regions of *D. melanogaster,* which lack *Drs* genes, are annotated below. Orthologous genes are shaded blue. A subset of gene symbols is shown to emphasise conserved gene arrangements. Grey annotations indicate non-syntenic genes. Pink annotations indicate lineage-specific genes. *Drosophila busckii* genes are annotated as gene ID and *D. melanogaster* orthologue (Table S2). **B)** The *D. melanogaster Drsl* gene clusters are located near each other on chromosome 3L (red). In *D. busckii,* most genes in this region are also located on chromosome 3L, indicated by annotations of orthologous BBH chromosome locations below *D. melanogaster* gene names. **C)** Phylogenetic cladogram and presence of *Drs* genes in Diptera. *Drosomycin* is restricted to only *D. busckii* and the Melanogaster group of *Drosophila.* A parsimony summary is given comparing emergence in a common ancestor followed by independent losses (Anc.) or assuming a single HGT event (HGT). Cladogram drawn from [48,51]. **D)** The *D. busckii* 2R *Drsl4* is immune-induced by heat-killed (HK) *M. luteus* injection similar to Toll-regulated *Bomanins (BomBc)*. Additional *Drsl* and *Dso* genes are shown in Figure S1.

#### The cluster 2R *Drosomycin-like* genes of *D. busckii* are immune-induced

We stimulated the immune response of adult flies by dipping a 0.1mm insect pin into heat-killed *E. coli* and/or *M. luteus* bacteria, then checked gene expression 24 hours post-injection. *Drosophila melanogaster Diptericin A (DptA)* is regulated by the Imd pathway and stimulated specifically by *E. coli,* while *Bomanin (Bom), Daisho (Dso)* and *Drs* are regulated by the Toll pathway and stimulated most specifically by *M. luteus* [8]. We recovered these patterns in *D. melanogaster* using our heat-killed bacterial preparations (Figure 1d, Figure S1). Surprisingly, we found that the *D. busckii DptA* orthologue, *DptC*, was strongly induced by both bacteria, and if anything, induction was higher following *M. luteus* pricking. However, *D. busckii Bom* and *Dso* genes were specifically induced by *M. luteus* and not *E. coli*, consistent with their expression pattern in *D. melanogaster*. By qPCR, we observed no inducibility of the *D. busckii* cluster 2L genes *DbDrsl1, DbDrsl2,* and *DbDrsl3* (Figure S1). However, both *DbDrsl4* (Figure 1d) and *DbDrsl5* (Figure S1) were induced specifically by heat-killed *M. luteus* injection, similar to the tested *Bomanin (BomBc).* At least for the Toll pathway, these gene expression patterns align with the specific presence of binding sites for the NF-κB protein Dorsal-related immunity factor (Dif) in *D. busckii DbDrsl4* and *DbDrsl5*, but not *DbDrsl1, DbDrsl2,* or *DbDrsl3* (Figure S2, purple arrows).

In summary, we confirm that among Diptera, the classical *Drs* locus is restricted to the Melanogaster group of the subgenus Sophophora. However, we unexpectedly recovered two *Drs* gene clusters on chromosome 2 in *D. busckii*. Considering genomic synteny analysis and the absence of *Drs* in other Diptera, the most parsimonious explanation for the presence of *Drs* in both *Drosophila* lineages is horizontal transfer of a *Drs* gene into the ancestor of *D. busckii*, the ancestor of the Melanogaster group, or possibly both stemming from an external source.

### *Drosomycin-like* genes are present in cockroaches and mantises

In a previous study [52], we built a profile HMM with the *D. melanogaster Drs* and *Drs-like* genes for use with HMMer-based searches and identified a *Drs* gene in the transcriptome of the oriental cockroach *Blatta orientalis*. Using this approach, we confirm the presence of 2 copies of *Drs* genes in the recently sequenced *B. orientalis* genome [53](Table S1). Notably, *Drs-like* genes were already found in the German cockroach *Blattella germanica* [52,54,55] but this genome was annotated as “contaminated” in GenBank at the time of writing, leaving the accuracy of those annotations unclear (assemblies GCA_003018175.1 and GCA_000762945.2, Data File S1). Here we have verified the presence of these *Drs-like* genes by PCR and Sanger sequencing and confirmed their presence in both *B. orientalis* and *B. germanica* (Table S3).

Upon confirming that the *Drs* genes of these *Blatta* and *Blattella* species were genuine, we scanned for the presence of *Drs-like* genes in other species of Blattodea, the order including cockroaches and termites (Table S1). Using newly sequenced genomes of 45 termites and 2 sub-social cockroaches [53,56], we confirmed previous results that *Drs* genes are absent in termite and *Cryptocercus* cockroach species [52,55] (Figure S3). However, using transcriptomic data [57] and other recently-available genome assemblies [58–61], we found *Drs* genes exist in most cockroach families (Blaberidae, Ectobiidae, Blattidae and Corydiidae, Figure 2). We further confirmed the presence of *Drs* genes in five additional species by PCR and Sanger sequencing (*Panchlora nivea*, *Blaptica dubia*, *Eublaberus serranus, Symploce pallens, Phoetalia pallida*) (Table S3). The breadth of species in which *Drs* is present strongly implies that *Drs* was also present in the ancestor of Blattodea. In the case of termites and related wood-eating *Cryptocercus* cockroaches, however, *Drs* has evidently been secondarily lost.

**Figure 2:**
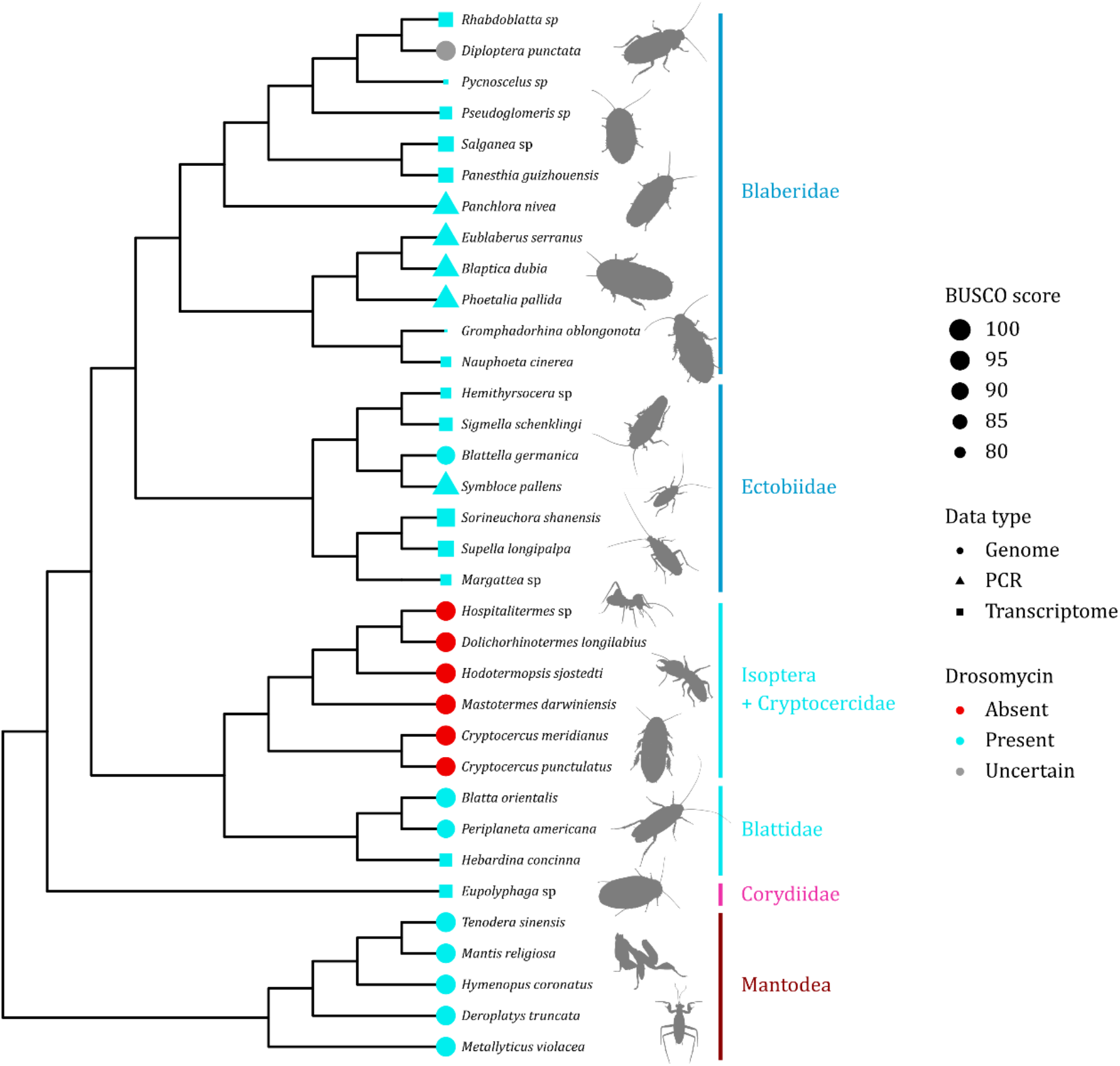
Phylogenetic cladogram of the Blattodea and Mantodea. The size of the branch tip is proportional to the BUSCO completeness score. Blue tips indicate that *Drs* gene(s) were found, red tips indicate that there is credible support for the absence of *Drs* genes (i.e. no hit found in multiple monophyletic genome assemblies). We conservatively annotate a lack of *Drs* recovered from *D. punctata* as “uncertain” (grey tip). Among Blaberidae this is the only assembly missing *Drs.* Independent validation with close relatives or additional specimens will clarify if this is genuine absence or an artefact of the specific genome assembly screened. *Drosomycin-like* genes were screened in genomic (circle tip), or transcriptomic (square tips) data or confirmed by PCR and Sanger sequencing (triangle tips). The abundance of positive hits in most Mantodea and Blattodea clades suggests that Drosomycin was acquired in the common ancestor and lost secondarily in Isoptera and Cryptocercidae. Cladogram based on [63–65]. Insect silhouettes from phylopics.org or designed from public domain images.

Having confirmed that *Drs* was present across Blattodea, we screened close outgroups from related insects using both tBLASTn and HMMer approaches. We confirmed *Drs* genes in five mantis genomes from the Mantidae, Hymenopodidae, Deroplatyidae, and Metallyticidae families, representing sublineages from key diversifications of the Mantodea order [62] (Figure 2). *Drosomycin* is therefore also broadly conserved in the Blattodea outgroup Mantodea.

Taken together, we infer that *Drs* genes in this lineage were present in the last common ancestor of mantises and cockroaches. *Drosomycin* was then secondarily lost in the ancestor of wood-feeding *Cryptocercus* cockroaches and termites. Overall, these results confirm and expand the annotation of the *Drs* gene family significantly beyond *Drosophila*.

### *Drosomycin* genes are scattered across the class Insecta

The presence of *Drs* genes in *Drosophila*, Mantodea, and Blattodea is surprising given the number of independent lineages between these insect orders thought to lack *Drs*. We therefore examined the extent of *Drs* presence/absence through a systematic screen across insects to better understand if *Drs* is unique to these lineages, or instead dispersed across insects more broadly; available sequence data was primarily sampled from winged insects (Pterygota). To this end, we initially screened up to 15 insect genomes from each unique branch of the insect phylogeny. When a *Drs* gene was found in two or more independent assemblies within a lineage, we annotated it confidently as “present”. If a *Drs* gene was absent in a clade where available genomic data was poor (i.e. assembly quality, availability of distinct species), we followed up using ad hoc searches with *D. melanogaster Drs* sequences, or a subset of *Drs* genes from the closest outgroup where *Drs* is found, to better annotate the presence and absence of *Drs* genes (e.g. tBLASTn against additional genomic assemblies, transcriptomic sequence read archives (SRAs)). In total, we screened >1400 genomes and transcriptomes from 25 winged insect orders and four outgroup orders: Zygentoma, Archaeognatha, Diplura, and Collembola (Table S1). Search methods included a HMMer-based HMM profile approach, as well as a recursive tBLASTn approach using a relaxed E-value threshold followed by manual curation [47,66]. An alignment showing a sampling of *Drs-like* genes we recovered is provided in Figure 3.

**Figure 3:**
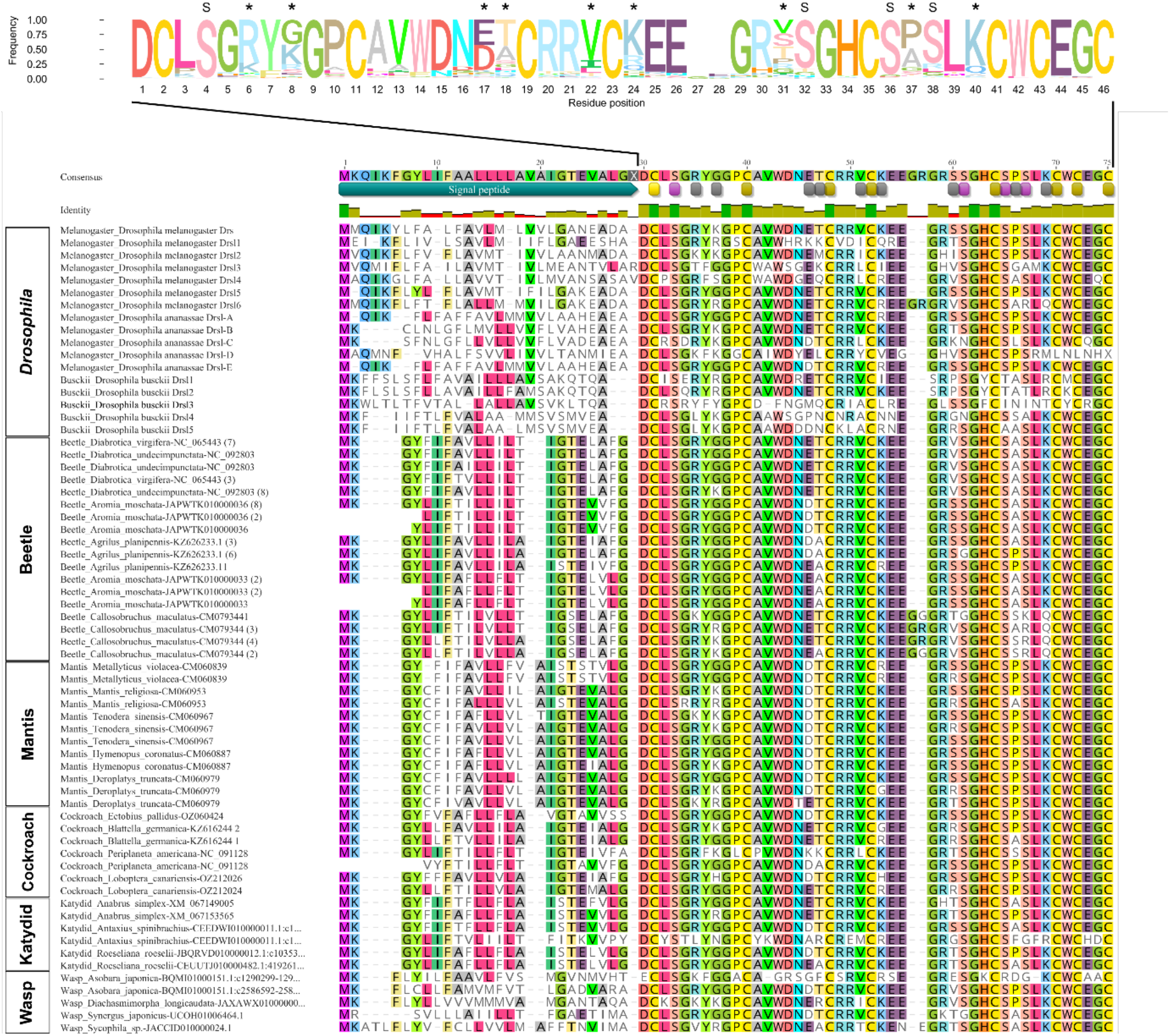
Representative full-length Drs peptides from diverse insects. Top: Sequence logo plot of mature Drs residues across discovered coding sequences. Polymorphic residues are highlighted by asterisks. Broadly-fixed serine residues are noted with “S” above their position. Bottom: Sampling of Drs full-length proteins. Highlighting indicates agreement to ≥65% similarity across all ungapped residues in the alignment. The alignment shown here is a subset of the total sequences contributing to (A). A list of recovered *Drs* genes and its mature peptide by HMMER search can be found in Data File S4 and Data File S5. Annotations under the consensus sequence note widely conserved serines (pink), cysteines (yellow), and polymorphic residues (grey).

#### Katydids have recently acquired Drosomycin

We first screened genomes and transcriptome SRAs from insect lineages closely related to Mantodea and Blattodea (Table S1) [67]. Here, we failed to detect *Drs* genes in any species of Phasmatodea (N=6), Embioptera (N=4), Grylloblattodea (N=2), Mantophasmatodea (N=4, but see supplementary text), Plecoptera (N=20), and Dermaptera (N=7). However, within Orthoptera (N=23), we did recover three species of Katydid (Tettigoniidae) with multiple *Drs-like* genes (*Anabrus simplex, Roeseliana roeselii, Antaxius spinibrachius*), including two *A. simplex* genomic assemblies derived from different samples (GCA_040419225.1, GCA_040414725.1). However, we found no other Orthoptera species with *Drs-like* genes, including the related Ensifera suborder species *Meconema thalassinum* (Tettigoniidae), *Myrmecophilus myrmecophilus* (Myrmecophilidae), *Gryllotalpa orientalis* (Gryllotalpidae), ten species of Gryllidae, and seven species of the suborder Caelifera (including the families Morabidae, Tetrigidae, and Acrididae).

Thus, we recovered *Drs-like* genes in three katydids, a broad diversity of mantises and cockroaches, but not in any other related insect clades representing three unique branches in the insect phylogeny. If *Drs* were acquired in the last common ancestor of Orthoptera, Mantodea, and Blattodea, explaining the presence of *Drs* in these lineages would require either a) four independent losses, including two within Ensifera (Orthoptera), one in Caelifera (Orthoptera), and one in the common ancestor of the insect orders Mantophasmatodea, Grylloblattodea, Embioptera, and Phasmatodea, or b) two gains: one within Tettigoniidae and one in the common ancestor of Mantodea and Blattodea. Under a simple parsimony model, the presence of *Drs* in katydids, mantises, and cockroaches, is better explained by two HGT events.

#### Drosomycin genes are present in two separate lineages of beetles

Extending our screen to >300 Coleoptera genomes, we discovered *Drs-like* genes in multiple beetle species. However, these genes were nearly exclusively present in two lineages. This includes the Elateriformia clade including Buprestoidea, Elateroidea, and Byrrhoidea. However, we note that while *Drs* genes were detected in all three species of Buprestoidea, only one out of 21 Elateroidea species and no *Drs* genes in the sole Byrrhoidea genome were identified. We also recovered a second clade of beetles with *Drs* genes including select Cucujiformia species from Chrysomeloidea, Cucujoidea and Curculionoidea. In the latter superfamily, we recovered *Drs-like* sequences in the Brentidae species *Aspidapion aeneum* and *Oxystoma pomonae* that appear to be pseudogenized; the *Drs* genes in these species contain indels that would cause loss-of-function, and we found no *Drs-like* genes in the other Brentidae species we screened (*Cylas formicarius*). We also failed to recover *Drs* genes in multiple Curculionoidea families including Attelabidae (*Apoderus coryli*), Anthribidae (*Platystomos albinus* and *Pseudeuparius sepicola*), and Curculionidae (15 genera). For other beetle lineages screened, we were unable to recover *Drs* genes including from Cucujoidea species (Nitidulidae, Kateretidae,), many Tenebrionoidea (but see supplementary text), Coccinelloidea, Cleroidea, the polyphaga clades Bostrichoidea, Scarabaeoidea, Staphylinoidea, Hydrophiloidea, Dascilloidea, and Scirtoidea, as well as outgroup species from Adephaga and Archostemata.

Thus, *Drs* is present in the distantly-related clades Elateriformia and Cucujiformia, but absent in at least ten other beetle lineages that branch from intermediate nodes between these clades [68]. It also seems that *Drs-like* genes were secondarily lost within the Curculionoidea, corroborated by the finding of putative pseudogenes in *Aspidapion* and *Oxystoma* – we suspect the *Drs* gene sequence has deteriorated in related Curculionoidea species too much to be positively identified.

Interestingly, the Drs protein sequences recovered from beetles bore a signal peptide more similar to cockroaches and mantises than to *Drosophila* both in sequence and in length (average percentage ID: Coleoptera vs. Dictyoptera (cockroaches and mantises) = 61.1%, and Coleoptera vs. Drosophilidae = 11.1%; DataFileS6; Figure 3b). Signal peptide regions typically experience relaxed selection, as the core of their function is to form hydrophobic alpha-helices, but the precise sequence that can accomplish this is highly malleable [69]. The high similarity of these signal peptide regions is unexpected, as the last common ancestor of holometabolan insects including beetles and flies was ∼333mya, and of cockroaches and holometabolan insects was ∼383mya [70]. Yet the striking similarity of *Drs* signal peptides from beetles and the cockroaches and mantises may suggest these genes share a more recent common ancestor.

#### Drosomycin pseudogenes are present in Liposcelis booklice

We also recovered four *Drs-like* sequences in the genomes of the barklouse *Liposcelis tricolor* (GCA_042257085), and one striking BLAST and HMMER hit (E-value < 1e-9) to what appears to be a *Drs-like* pseudogene in *Liposcelis bostrychophila* (GCA_037577465). However, we found no *Drs-like* sequences from *Liposcelis brunnea* (GCA_023512825), or four other Psocodea species (*Columbicola columbae, Loensia variegata, Mesopsocus fuscifrons, Psococerastis gibbosa*), even using the *Liposcelis spp.* sequences as a query. Assessing the *L. tricolor* putative *Drs-like* loci, each contained loss-of-function mutations (e.g. premature stop codons, loss of start codons). Still, for one *Drs* of *L. tricolor,* an open reading frame lacking a signal peptide was recovered that could feasibly encode a full-length Drs peptide, though it would lack a secretion mechanism (hereafter *Ltri\Drsl-ψ1*). We therefore checked a transcriptome SRA of *L. tricolor* to determine if this sequence was expressed (SRX25805314), however no reads matching *Ltri\Drsl-ψ1* were recovered. Given *Drs* occurrence in *L. bostrychophila* and *L. tricolor,* but not *L. brunnea,* we argue that a very recent horizontal transfer has occurred, which introduced *Drs* into the common ancestor of *L. tricolor* and *L. bostrychophila*. However, *Drs* is not ancestral to Psocodea, and was rapidly pseudogenized after its introduction into an ancestral *Liposcelis* species (Figure S4).

#### Drosomycin is found in three parasitic wasp lineages

In a screen of >700 Hymenopteran genomes, we found *Drs* genes present in three unique lineages of parasitic wasps (Braconidae, Chalcidoidea, and Cynipoidea). Both the Chalcidoidea and Cynipoidea are part of the infraorder Proctotrupomorpha, and the sequence similarities within Braconidae was higher than in these lineages (Braconidae = 85.3%, Chalcidoidea = 73.7%, and Cynipoidea = 73.5%). We were unable to recover any *Drs* genes in two other poorly sampled lineages of Proctotrupomorpha (Platygastroidea and Diapriodea) and many species of Chalcidoidea and Cynipoidea, suggesting a potential lineage specific loss of the *Drosomycin* in these families. Despite large sampling (>130), we also did not recover any Drs genes in Ichneumonidae, the sister family of Braconidae. This finding would require either one gain in the ancestor of Hymenoptera and 6-8 losses, or 2 gains (one in Braconidae and one in Proctotrupomorpha) and 2 losses. Thus, the most parsimonious explanation for *Drs* distribution within these parasitic wasp lineages is that these Drosomycin genes were acquired by the Proctotrupomorpha and Braconidae ancestors independently.

#### The presence of Drosomycin in insects is relatively rare

We did not recover *Drs-like* sequence in any species from the insect orders Odonata, Dermaptera, Plecoptera, Grylloblattodea, Mantophasmatodea, Embioptera, Phasmatodea, Thysanoptera, Hemiptera, Raphidioptera, Megaloptera, Neuroptera, Strepsiptera, Trichoptera, Lepidoptera, Siphonaptera, and Mecoptera. The distribution of these insect lineages, alongside the presence of *Drs-like* genes and pseudogenes we have described, makes an insect-ancestral *Drs* locus unlikely under an evolutionary model based on parsimony. Indeed, to explain the results of our screen, presuming a shared ancestry from within class Insecta would require 64 independent loss events since those lineages diverged from a putative *Drs*-encoding last common ancestor (Figure S5). While we observed *Drs* pseudogenes in *Liposcelis* species and found good evidence of gene loss in *Cryptocercus* cockroaches and Isoptera, as well as a subset of Curculionoidea, such a systematic loss of Drosomycin in these 64 lineages, (with conservation in just nine, excepting booklice), would represent a remarkably specific pattern of evolutionary retention and loss. Alternatively, presuming all *Drs* gene lineages in insects stem from horizontal transfer events, only nine HGT events are required to explain the full diversity of insect *Drs* genes and pseudogenes.

To estimate the likelihood of these *Drs* genes as the product of HGT events, we reconciliated the species tree of all insects screened (Data File S2) with the *Drosomycin* gene tree (restricting our selection to one species per genera or *Drosophila* group). We produced a hundred gene trees of the *Drosomycin* gene using Bayesian phylogenetic analysis implemented in MrBayes and reconciliated each tree with our insect phylogeny. We calculated the number of unidirectional HGT events inferred per lineage pairs across the 100 reconciliated gene trees and used the average over four replicate analyses as a measure of support for that event (Data File S3). Among the HGT events with at least 70% support, five events were inferred across orders (Braconidae wasp-*D. busckii*: 78.5%; Braconidae wasp-*D. melanogaster*: 70.5%; Katydid-Proctotrupomorpha wasps: 94.5%; Mantodea-Katydid: 75%; Cucujiformia beetles – Blaberid cockroaches: 70.25%) and six within orders (Blattodea, Hymenoptera, Coleoptera). Other HGTs with lower confidence on the origin was inferred among Mantodea, Coleoptera, Blattodea, Orthoptera, and Proctotrupomorpha wasps, and a single event between Braconidae wasp and Curculionoidea beetles (10.75%).

Collectively, these results support the hypothesis that at least one HGT event has occurred across insects, and likely more than one. The proposed HGT events between Braconidae wasps and *Drosophila* species are intriguing as we recovered *Drs* genes in *Asobara* wasps known to parasitize *Drosophila*, a process that offers an intuitive mechanistic process with potential for HGT. While other HGT events were also proposed by this analysis (including a consistently high support for Proctotrupomorpha plant parasitic wasps and katydids), a potential mechanism is less clear.

### *Drosomycin* similarity is greater than expected of an ancestral conserved gene

To rule out the possibility that our search approaches are insufficiently sensitive to recover *Drs* across insects, we next performed a similar sequence search for a widely conserved gene. We chose the Dorsal NF-κB transcription factor, a gene with roles both in dorsoventral patterning and immunity [71]. Specifically, we used the IPT domain of Dorsal, with a query approximately size-matched to Drosomycin, and with a similar degree of conserved residues, to achieve comparable search sensitivity for HMMer and BLAST screens across species. Using the Dorsal IPT domain protein sequence, we could recover Dorsal orthologues in all species screened using HMMer and our recursive tBLAStn approach (Table S1). Thus, our search algorithms are sufficiently sensitive that we would expect to have recovered *Drs* in most insects if it were ancestrally present and universally conserved.

To test if Drs protein sequence is more similar across species clades than would be expected by common ancestry, we compared the percent similarity of protein alignments of Drs and the IPT domain within and between clades of insects. Within clades, we found that the percentage identity of IPT_Dorsal was not higher than for the mature peptide sequence of Drosomycin (Mantel-Haenszel chi-squared test, χ2 = 19.1, p > 0.999). However, the percentage identity of the consensus sequence among clades was significantly higher for Drosomycin compared to IPT_NF-κB (Mantel-Haenszel chi-squared test, χ2 = 103, p < 0.001). This indicates that mature Drs peptides are more similar across distantly related species than expected, when compared to a conserved region of an essential development/immunity transcription factor.

We further found that many Drs residues are extremely well-conserved across ≥75% of sequences. However, we also identified multiple polymorphic sites where alternate residues are common (Figure 3). While it is beyond the scope of this study to functionally test the consequences of these many polymorphisms, we highlight these residues of potential interest for future studies investigating Drs-like proteins.

### Codon usage suggests a common origin of *Drosomycin* in disparate insects

If *Drs* genes have evolved convergently, we may expect to see certain common protein residues produced via alternate codons. To investigate this possibility, we extracted the full-length *Drs* coding sequences from Elateroidea and Cucujiformia (Coleoptera), Braconidae and Proctotrupomorpha (Hymenoptera), Mantodea, Blattodea, Orthoptera, and *Drosophila,* and using codon alignments, performed sequence and phylogenetic analyses. Given the immense timescales involved, which allow silent mutations to fluctuate forwards and backwards, we restricted our analysis to codon usage of conserved serine residues. Serine is unique among amino acids as eukaryotes encode serine either via TCN or AGY codons (N = any nucleotide, Y = C or T). Simple transitions between one serine codon type and the other are rare due to the need for an intermediate non-synonymous mutation event. As a result, essential serine codons often become fixed [72]. There are four conserved serine residues across the *Drs* genes of flies, beetles, mantises, katydids, wasps, and cockroaches (Figure 3). To overcome the challenge of copy number variation across species, we used the modal codon (i.e. most common codon) for each species’ *Drs* sequences at each serine site as representative of that species, and performed our analysis with each unique species as an independent sample within their respective clades.

Serine 1 is universally encoded by a TCN codon (100.0% TCN, Figure 4a). In all lineages except for wasps, the profile of the next three serines shows a likely ancestry utilising AGY codons at each site; in the case of *D. busckii,* the modal serine 3 codon shows a pattern consistent with either of the TCN or AGY codon being the likely ancestral codon. We next used a Bayes Factor (BF) approach to compare the probability of a pattern arising from homology or convergence, with interpretation provided along the Jeffreys’ scale [73], where BF > Log10(100) (i.e. +2) suggests homology, while BF < Log10(0.01) (i.e. -2) suggests convergence. Across comparisons, the Melanogaster, Elateroidea, Cucujiformia, Mantodea and Blattodea, and Katydid *Drs* lineages share serine codon usage strongly suggesting convergence (comparisons among these lineages are consistently log10(BF) > +2, and often log10(BF) > +3). Due to low sample size (n = 1), *D. busckii* cannot be assessed statistically, but consistently trends towards homology with these clades. However, comparisons of these lineages to the wasp lineage Braconidae resulted in broadly neutral Bayes Factor values ( -2 < log10(BF) < +2), with the exception of comparison to Cucujiformia likely owing to mixed codon usage in serine 2. On the other hand, comparisons to Proctotrupomorpha consistently resulted in negative log10(BF) values, often < -2, with the greatest difference being comparison to Braconidae (log10(BF) = -7.5. This pattern is driven by usage of an AGY codon at serine 3 in this lineage, not found in any other multiply-sampled lineage.

**Figure 4:**
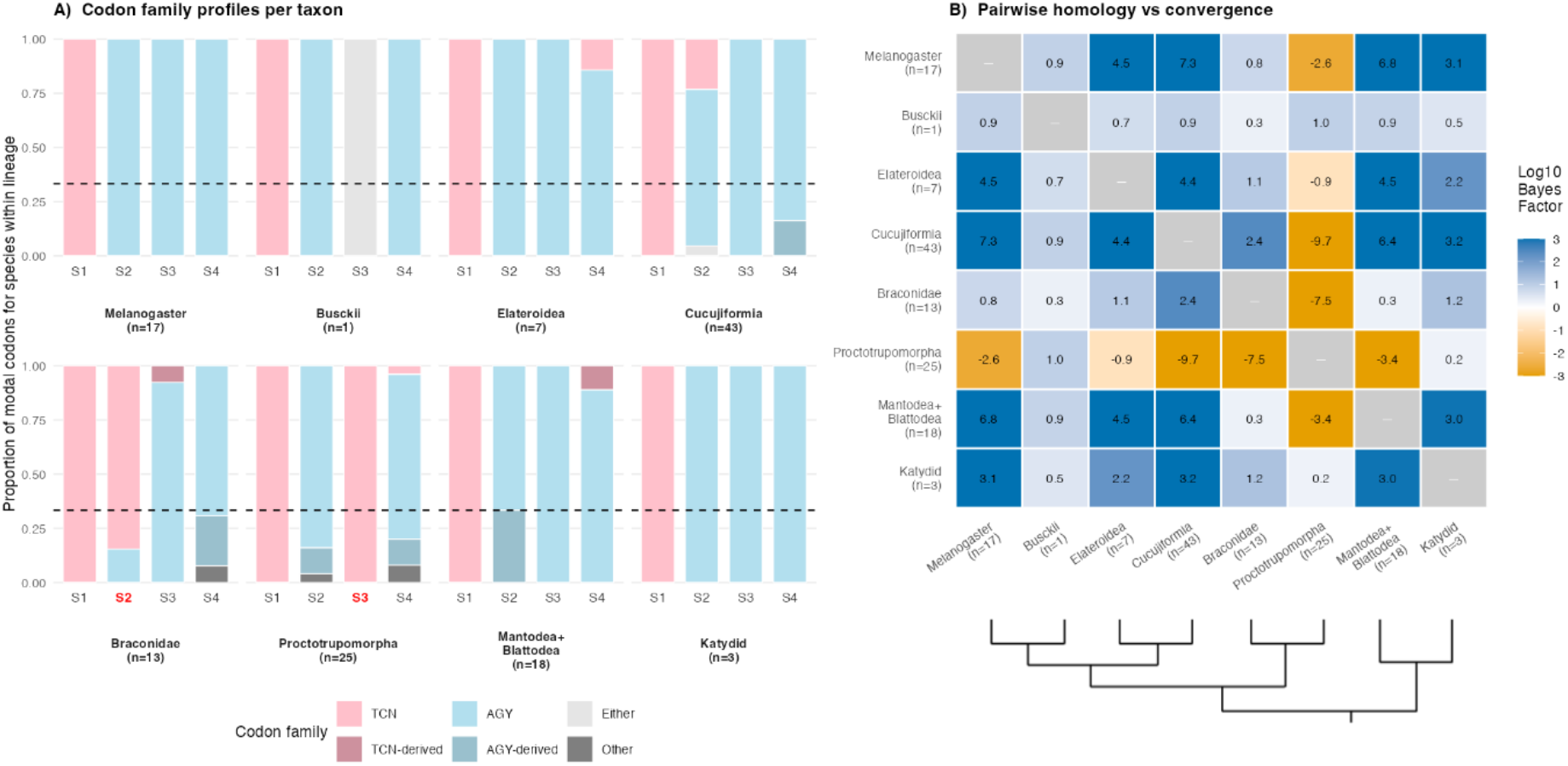
Serine codon analyses indicate broadly homologous sequence usage across disparate insect taxa. A) Serine codon usage in flies, mantises and cockroaches, katydids, and beetles all follow a similar S1 = TCN, S2-S4 = AGY codon usage pattern. On the other hand, two lineages within Hymenoptera show divergent serine codon usage with Braconidae species using TCN at S2 and Proctotrupomorpha species (including Cynipidae and Chalcidoidea) use TCN at S3 (highlighted in red). The dashed line indicates the expected proportion (1/3) of AGY codons under a random convergence scenario. B) Jeffreys’ scale Bayes Factors (BF) for pairwise comparisons of each lineage assessing the probability of independently deriving the same serine codon usage pattern across four codons. Blue indicates likely homology of sequence and yellow indicating likely convergence of sequence. A log10 BF > 2, or -2 > BF, is considered “decisive” evidence in Jeffreys’ scale. All non-Hymenoptera lineages sharing the same S1-S4 codon usage patterns have a high BF in pairwise comparisons, suggesting homology. However, comparisons between wasp lineages to each other, or to other lineages, are either inconclusive (Braconidae), or broadly suggest convergent evolution underlies serine codon usages relative to other lineages (Proctotrupomorpha).

Taken together, Bayes Factor analysis consistently suggests homology among these groups. However, the wasp *Drs* genes show mixed signals in terms of a direct homology, and utilisation of a TCN codon at serine three in Proctotrupomorpha suggests this lineage has evolved that serine codon convergently. Taken alongside species tree and gene tree reconciliation analysis (Data File S3), *Drs* genes of Proctotrupomorpha may stem from an independent genetic source compared to other insects. On the other hand, serine codon analysis suggests the mature *Drs* coding sequence of flies, beetles, cockroaches, mantises, and katydids is homologous.

Finally, we performed phylogenetic analyses of *Drs* lineages to determine if *Drs* coding sequence would support monophyletic or paraphyletic clustering. Due to the short length of this gene, being ∼130 nucleotides ungapped if including the signal peptide and mature peptide, results from phylogenetic analysis are not expected to be robust, but can nonetheless inform on the direction of the trend. Consistent with horizontal transfer, multiple lineages clustered paraphyletically in MrBayes consensus trees utilising either the full-length coding sequence or just the conserved Drs peptide coding sequence, with wasp and aly lineages clustering together forming long branches relative to other lineages (Figure S6). Moreover, various *Drs* genes clustered paraphyletically between e.g., cockroaches and beetles, or katydids and wasps, indicating that this gene sequence is insufficiently diverged between these lineages to faithfully recapitulate the true phylogeny, suggesting a more recent common ancestry of *Drs* genes across insect lineages compared to the phylogenetic species tree.

Taken together, *Drs* presence in disparate insect lineages is difficult to explain through shared ancestry and subsequent loss. Phylogenetic analysis shows paraphyletic clustering, and sequence similarity analysis compared to the Dorsal IPT domain suggests *Drs* sequences across lineages share a more recent common ancestor than the true insect phylogeny. Serine codon usage further supports a homology of *Drs* sequence across disparate insect lineages last sharing a common ancestor ∼383 mya, but also reveals a striking divergence of codon usage in *Drs* genes of parasitic wasps, particularly Proctotrupomorpha plant gall wasps; this result suggests at least two independent sources of the *Drs* genes within insects, or perhaps rapid divergence within the wasp lineages after *Drs* introduction from a common source. Parsimony analysis is also highly incongruous with a *Drs* gene presence in the common ancestor of winged insects followed by subsequent losses, while proposing HGT into various clades greatly reduces the total events required in parsimony analysis. In summary, these findings strongly suggest that *Drs* has been horizontally transferred across insect lineages at least once, and likely multiple times.

## Discussion

The distribution of similar AMPs across animal taxa has long posed challenges to understanding AMP sequence conservation and diversification. AMPs evolve rapidly, including examples of both diversifying and balancing selection [9,10,74,75]. Immune effectors including AMPs can also vary in copy number between individuals within a population, which can even become fixed in species, possibly explaining increases in host susceptibility to parasites and pathogens [9,16,47,76]. Here we reveal the potential for the reverse scenario: the gain of an AMP equipping a host immune arsenal with a novel antifungal peptide gene.

Recent studies, particularly in *Drosophila,* have emphasised the importance of AMP polymorphisms in determining defence competence against specific pathogens in vivo [9,10,16,40,77]. In the present study, we have not validated the antimicrobial activity of any of the recovered *Drs* peptides. However, there is a longstanding literature showing antifungal activity of *Drs* and *Drs-like* Defensin peptides both in vitro and in vivo [43,46,78–80], including ones that we have highlighted. It is likely that at least some of the novel *Drs* sequences encode peptides with activity against fungal infections. A fruitful future endeavour could be to focus on the impact of common polymorphisms on *Drs* antimicrobial activity. Equally, *Drs* in *Drosophila melanogaster* has been associated with anti-tumour activity [81,82], response to traumatic brain injury [83], and *Drs-like* genes are expressed in the *Drosophila* midgut and other surface epithelia under the regulation of JAK-STAT and NF-κB signalling [84–86]. *Drosomycin* can further interact with sodium-gated ion channels similar to scorpion neurotoxins [87], consistent with the potential to interact with host neurology and physiology. Moreover, plant defensin-like peptides have been shown to regulate pollen tube development, including by interacting with potassium channels [88,89]. The reason *Drs* has such pluripotent functional potential could stem from e.g., an ancestral role for *Drs* as a bacterial or viral toxin that acted as a virulence factor to disrupt eukaryote neurological ion channels, which has now been domesticated. Indeed, recent studies found that flies, aphids, and wasps have domesticated horizontally-acquired bacterial toxins to serve as immune defences, or toxins contributing to parasite attack [90–92]. In this regard, an intriguing possibility is that the Proctotrupomorpha gall wasps have divergent *Drs* genes not solely because of convergent evolution, but also because *Drs* has become adapted in these parasitic wasps to manipulate plant hosts, potentially acting with different focal roles as venom peptides compared to the immune or physiological roles of domesticated insect *Drs* in host physiology.

Here we propose that *Drs* has been horizontally transferred into different insects at least once, and likely more than once (Figure 5). A scenario supported by our reconciliation of the *Drs* gene tree with our species tree suggests one or more HGT events between braconid parasitoid wasps and *Drosophila*. Previous studies have shown parasites can act as vectors of symbionts [93], and that parasitoid wasps inject virus-like particles to manipulate host biology [94], which provides for an intuitive mechanism enabling *Drs* HGT through host DNA damage incurred during the stress of infection. It may even be that parasitic wasp *Drs* genes are co-opted by the parasite genome for host manipulation, an intriguing speculation that merits future study.

**Figure 5:**
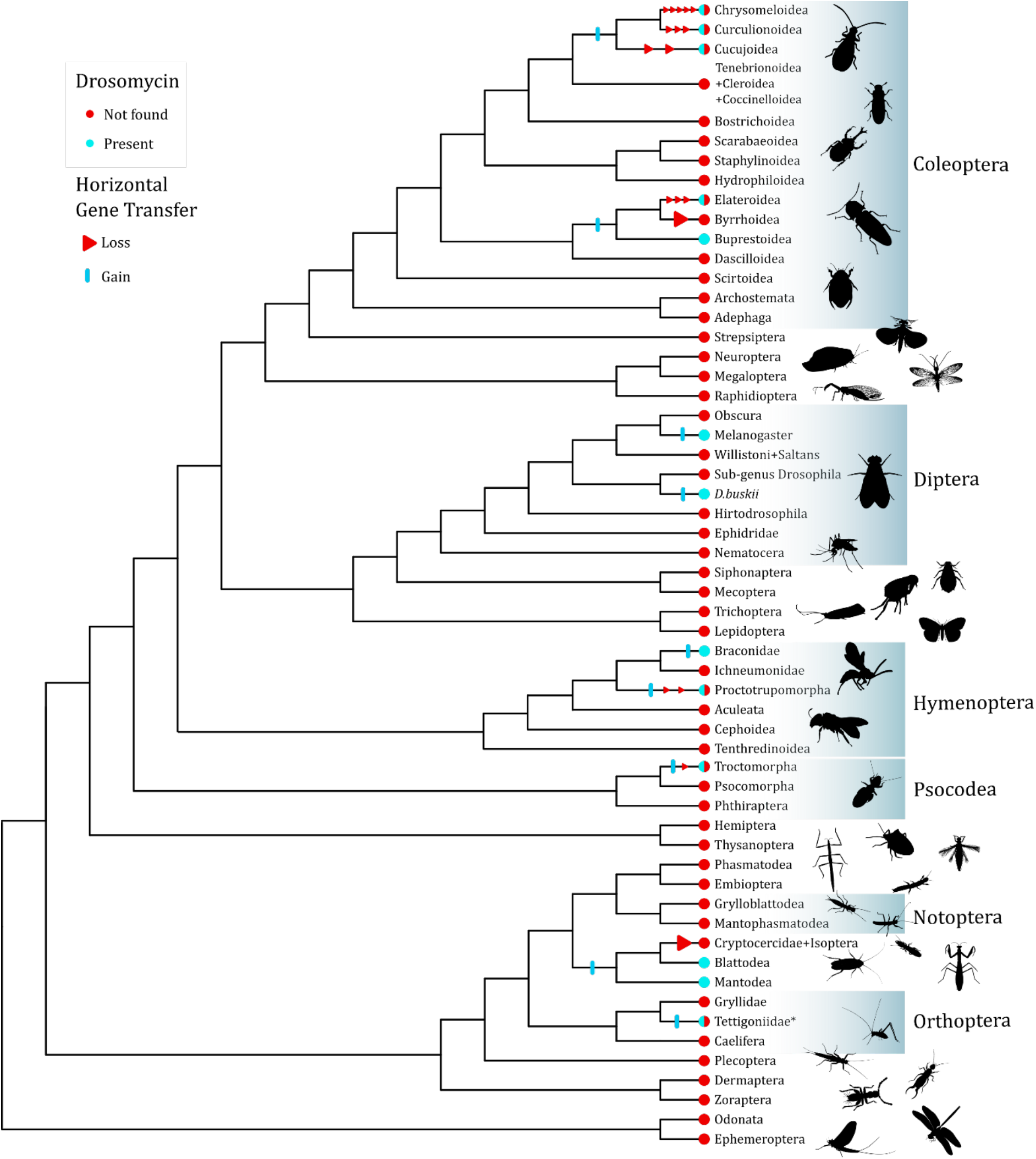
Simplified overview of the presence and absence of *Drs* genes across relevant insect clades. In multiple places, independent lineages are omitted to simplify the figure. Blue tips indicate lineages where *Drs* genes (including pseudogenes) were found, while red tips indicate the many other lineages where we failed to recover *Drs* genes. Half-filled circles indicate both presence and absence in at least some lineages. While the most parsimonious explanation for this pattern of gene evolution would involve 20 gain and loss events, we consider some secondary loss events highly plausible in some lineage and therefore, we favour a scenario with nine gain events (blue bars along branches) and 18 secondary loss events (red triangles along branches). A summary of parsimony counts for the common ancestor hypothesis are presented in Figure S5. Note that within the Tettigoniidae (*), the gain of *Drs* occurs in a monophyletic sublineage. Insect silhouettes are from phylopics.org.

Our reconciliation analysis also proposes multiple transfers among katydids, beetles, mantises, cockroaches and Proctotrupomorpha wasps. While some of these events are possible, some proposed transfer events include transfers between e.g. two Blaberidae species, which is not required to explain presence in both species. The very high support for a Proctotrupomorpha and katydid HGT event is intriguing, but unlike braconid wasps, Cynipoidea and Chalcidoidea wasps are plant gall parasites and do not parasitize katydids. It may be that the katydid result (n = 3 lineages) stems from a low sample size, or could reflect an intermediate host plant (i.e. wasp <-> plant <-> katydid).

A single common ancestor of all insect *Drs* genes diversifying with the species tree would require the retention of *Drs* in various extant lineages, but >60 independent losses in related lineages; a complete parsimony account for this idea is outlined in Figure S5. While we observed patterns likely explained by ancestral presence and secondary loss (e.g., *Cryptocercus spp.*, various cucujiform beetle species) and could even recover *Drs* pseudogenes in some lineages (e.g., *Liposcelis spp.,* Brentidae beetles), it would be highly improbable that all intermediate lineages uniformly lost *Drs*, leaving not even traceable pseudogenes across well-sampled lineages such as the genomes of most Diptera. On the other hand, a strict parsimony argument would propose 20 events (18 gains + 2 losses or 14 gains + 6 losses) to explain *Drs* patterns across insects. However, considering secondary losses to be likely in different lineages, we believe that a more plausible pattern would be 9 gains and 18 losses (gains/losses summarised in Figure 5, and see Figure S5). While other patterns of gain/loss are possible, our interpretation depends on available genomic data and phylogenetic accuracy (ex. the beetle phylogeny is not settled [95]). It is likely that future sequencing efforts and additional screening would recover additional *Drs* genes that we have missed; indeed, we found *Drs* sequences in two Collembola species (hexapods, but not insects) and a single species of Tenebrionidae (*Lagria hirta*), which we have not included in our summaries due to insufficient sampling for confidence in reporting (Data File S6 and see supplementary text). Importantly, patterns of genomic synteny (Figure 1), high sequence similarity (Figure 3), uniform serine codon usage across disparate clades (Figure 4), paraphyletic sorting of lineages (Figure S6), and reconciliation analyses, each suggest a more diverse evolutionary history compared to a highly non-parsimonious presence of *Drs* in the common ancestor of most winged insects (Figure S5).

The horizontal transfer of *Drs* genes into other lineages is not necessarily expected to result in a novel immune-inducible gene. Indeed, we found pseudogenization of *Drs* sequences in two *Liposcelis* booklice that seem evolutionarily young. Yet in *D. busckii* we observed immune inducibility of *Drs* genes (Figure 1d). The regulation of NF-κB genes is known to rely greatly on the presence of NF-κB motifs within 200bp of the transcription start site [96–98]. It is tempting to speculate that vectors of HGT, such as transposons activated or transmitted during the stress of parasitisation, could therefore insert short single-exon AMPs like *Drs* with core regulatory elements into novel genomes as a pre-existing promoter + gene package: for instance, the total length of the 200bp of *Drs* promoter and complete transcribed region is just 587bp. Thus, the retention of horizontally transferred immune peptide genes may be aided by their key promoter elements being common across the tree of life [6] and found in close proximity, making co-transfer likely. Indeed, a recent study in plants showed horizontal transfer events are more likely to be retained when self-regulating [99].

Imagining *Drs* was horizontally transferred into various insect lineages, such promiscuity could result from transposable element (TE) activity, known to underlie HGTs of other animals [100]. Indeed, within AMP lineages, tandem duplication events are common, and we recovered similar events in *Drs* gene loci across insects (e.g. Figure 1ab, and see [44]). However, in an initial screen, we did not find evidence for a common set of TEs in the vicinity of *Drs* genes that could explain their presence across insects (Figure S7, Table S4, and see supplementary text). The introduction and duplication of *Drs* genes in certain lineages could result from clade-specific mechanisms, or alternatively, historic lateral transfer events where signatures of HGT have been lost over deep evolutionary time.

We also observed pseudogenization and loss events in multiple insects (*Cryptocercus* spp., curculionid beetles, *Liposcelis* spp.). Previous studies have suggested that diet shapes the host microflora, and that AMPs influence host epithelial and gut microbiota communities [5,11,101–103]. It would be interesting to learn if there was a meaningful dietary shift in the ancestors of various beetle species where our screen might suggest repeated losses. Indeed, *Cryptocercus* spp. cockroaches and termites represent a clear case of *Drs* gene loss coinciding with adaptation to a novel wood-feeding diet [53,56]. This finding parallels a recent evolutionary study of the fruit aly *Diptericin B* gene, an AMP with specific potency for controlling *Acetobacter* bacteria that are commonly found in rotting fruit. That study found that *Diptericin B* is secondarily lost in hosts that diverged to feed on plant tissue or mushrooms where *Acetobacter* is absent [16]. Sociality itself could play a role in shifting pathogen pressures, as termites may have relatively weaker responses to infection than cockroaches [52], and social ants were recently shown to lack an immune response to fungi, perhaps compensated for by social grooming (i.e. behavioural immunity) [104].

Alternatively, AMPs may impose a fitness cost on host species, resulting in hosts navigating trade-offs between e.g., improved defence or negative fitness effects, such as microbiota or endosymbiont dysregulation [23,26,27,105–107]. The immune response can also induce autotoxic damage, and certain AMPs, including polymorphisms, may reduce reproductive success [40,41,108]. In vertebrates, Defensin-like AMPs can also act as signalling molecules, interacting with host receptors to modulate immune effector cells [101], or even to determine fur colour [109]. Thus, the horizontal transfer of AMPs could impact hosts in unexpected ways, even if their sequence is originally adapted for microbicidal activity.

While we have focused on *Drs*, the same evolutionary principles described here could apply to other peptides. For instance, widespread proline-rich peptides, such as Drosocin, Abaecin, and Pyrrhocoricin, commonly encode Tyrosine near the N-terminus required for ribosome binding, and an O-glycosylated threonine within a PXPT motif that is critical for antibacterial activity [21,110,111]. While these features could reflect convergent evolution towards a successful antimicrobial strategy, ancient horizontal transfer events could explain this motif across proline-rich peptides of diverse lineages. While it was originally proposed that *Drosocin* of *D. melanogaster* emerged from a tandem duplication of an ancestral Attacin gene [17], *Drosocin* orthologues were later found seemingly precisely excised and translocated into the gene region of *Gr43a* in other flies [74]. Given the evidence supporting *Drs* HGT shown here, it may be that *Drosocin* was in fact horizontally transferred into one or both loci, avoiding the need to explain both loci through precise and complex genomic rearrangements. As future studies continue to investigate AMP histories and AMP-microbe interactions, it will be interesting to consider both ecological and evolutionary factors that could explain their gain, retention, and loss.

## Conclusion

Here we have outlined numerous lines of evidence to support at least nine HGT events of the antifungal peptide gene *Drosomycin* across insects. Understanding the origins of AMPs is difficult over long evolutionary timescales owing to their short sequence and the technical limits of search sensitivity. Our study can be considered alongside the developing literature on the domestication of bacteria and their toxins for host defence [90,91,112] and parasite attack [92], suggesting the possibility that *Drs* could have been horizontally acquired from a source microbial species not present in sequence databases. Our study also contributes to a broad body of literature on the rapid evolution of host immune genes, and comes at a time of increasing appreciation for how AMP variation can mediate infection susceptibility. Indeed, if AMP polymorphisms and losses can be major determinants of infection outcome across animals [11], the gain of an AMP in a host’s immune arsenal could dramatically alter infection dynamics in the same way. Given recent and ongoing zoonotic pandemics caused by bacterial, fungal, and viral pathogens, exploring the complement of AMP genes across species may reveal lineage-specific effectors that help explain differences in defence competence across species. Why might such lineage-specific effectors exist? Here we show that ait-for-purpose AMP genes can not only be lost between related species, but they can also be gained just as suddenly.

## Material and Methods

### *Drosomycin* search strategy with tBLASTn

Because *Drosophila* species have many high-quality annotated genomes, we used, as in a previous study [47], a reciprocal tBLASTn approach to screen Drosophilidae species for *Drs* presence inputting just the mature *Drs* peptides of *D. melanogaster* (and subsequently *D. busckii*) as queries. Due to the short query sequence of just ∼44 mature peptide residues, we used an artificially high E-value (E > 1000) as a cutoff for returning BLAST hits followed by a manual curation of the best blast hit. This returns many sequences of dubious similarity but ensures true hits are not discarded due to the limits of the short query length. This relaxed E-value is required as the length of the mature peptide of 44 residues restricts the minimum possible E-value to just 7.50e-25 when using the *D. melanogaster* Drs protein to tBLASTn against the Dmel_R6 reference genome (standard search parameters) (Figure S8). Using a modified Drs sequence substituting at every 3rd residue results in a tBLASTn search result of anywhere from 3.6e-14 to just 2.0e-6 depending on which residues are substituted, despite each of these artificially modified Drs protein sequences being ∼68% identical to the original *D. melanogaster* Drosomycin. Further disruptions cause rapid dropout of the E-value, leaving true positives with identifiable homology indistinguishable from false positives; thus our approach to use a high E-value alongside manual curation for BLAST-based searches. In total, we screened all species of Drosophilidae available on GenBank, alongside a manual curation effort including 84 species of Drosophilidae and 46 species of outgroup Diptera (Table S1, Data File S7). The same procedures were used in batch searches to screen *Drs* genes across insects. In addition to queries using the full diversity of *D. melanogaster Drs* and *Drs-like* genes, we also used a selection of *Drs* genes from the closest outgroup to the species being searched to give the best chance of recovering related genes or pseudogenes. BLAST searches were conducted using a local BLAST database searching one assembly at a time in Geneious Prime 2025.2.2, and tBLASTn was carried out with BLAST package version 2.12.0.

### *Drosomycin* search strategy with HMMER

Genomes and transcriptomes analysed in this study (Table S1) were also screened using HMMer. To do so, genomes and transcriptomes were first six-frame translated using a customed Python script (https://github.com/CedricAumont/DrosomycinHGT). All possible coding sequences longer than 15 amino acids were kept and saved in fasta format. Information on the sequence (contig number, strand sense, genomic position) was stored. A *Drs* HMM profile was built using HMMER version 3.3.2 (https://www.hmmer.org/) and a MSA of all *Drs* sequences from *Drosophila melanogaster* downloaded from OrthoDB (www.orthodb.org) using MAFFT (v7.475) with the G-INS-i option and 1000 iterations [113] and reformatted into Stockholm format with the esl-format command from HMMER. The *Drs* profile HMM was used to search all six-frame translated proteomes. We used the domain search option from HMMsearch and filtered out all hits in which E-values for full sequence and inclusive domain were above 1e-3. Hits were given a unique ID incorporating information on its length, strand sense, scaffold number, alignment position on the profile HMM, and genomic position. The hits were sorted by scaffold and strand and ranked by genomic position within each scaffold. If necessary, partial hits were reconstructed automatically if their parts aligned contiguously on the HMM profile and were from the same strand sense and same scaffold number. Sequences that covered more than 75% of the profile HMM were stored in fasta format.

### Validation of search strategies

As failure to find a gene in a species might be due to the sensitivity of our search strategies, we tested our method with a similarly sized region of a conserved and intronless domain against a subset of the total genomes we screened (Table S1). This included 132 genomes screened by tBLASTn and 169 genomes screened by HMMer; numbers of assemblies screened differ due to differences in local database and pipeline setups. We targeted the IPT domain of the *dorsal* NF-κB gene.

For the BLAST search, we initially used the *D. melanogaster* segment as a tBLASTn query. In 125 cases the *dorsal* gene was recovered with e-value ≤ 1e-5. In two cases, the best blast hit was the insect’s *dorsal* gene, but with e-value > 1e-5. In the remaining cases (all Lepidoptera species), we moved on to using a *dorsal* gene segment recovered from *Bombyx mori* via our initial *D. melanogaster* search, which retrieved dorsal in the remaining lepidopteran species. Thus our recursive and iterative BLAST approach succeeded to recover *dorsal* in all species screened, including insects both with and without *Drs*.

Separately, we independently screened 169 genomes with our HMMER-based approach. We aligned the *dorsal* and *Dif* genes from *D. melanogaster* and built the HMM profile with the same protein domain region. With the HMMER-based search we recovered the IPT domain in all but two genomes. One absentee was due to the low quality of the genome (BUSCO completeness of 34%, *Celatoblatta quinquemaculata,* GCA_048987405.1) and the second genome was fragmented in the genomic region of the domain and so the hits found did not cover contiguously at least 75% of the HMM profile (*Polylopha cassiicola*, GCA_038024825.1). The later was successfully recovered by the BLAST searched using the *Bombyx mori Drs* as query. Therefore, we confirmed that our HMMER-based search strategy recovered the targeted gene in all species of sufficient genome quality and completeness and that our methods are complementary into recovery all sequences.

### Cockroach *Drs* presence screen

PCR primers were designed to confirm the presence of *Drs* in some species of cockroaches. The primer design was based on sequences found during the screening of the genome and transcriptome of *Blaptica dubia*, *B. germanica*, and *Panchlora nivea*. Primers were designed using the NCBI primer design tool (https://www.ncbi.nlm.nih.gov/tools/primer-blast/) and followed by manual selection (Data File S8).

The cockroaches were collected from their rearing boxes and euthanized via freezing 30 minutes prior to DNA extraction. The head of two cockroaches per species was used as input material. All DNA extractions were carried out with the DNeasy Blood & Tissue Kit (Qiagen, Germany) following manufacturer instructions. Briefly, heads were cleaned with 99% molecular-grade ethanol, dried, and homogenized with PBS in a tissue ruptor (MP Fastprep-24 5G) with one 2-mm glass bead. Tissue were incubated in lysing buffer with proteinase k at 56°C for 10 minutes. To avoid RNA contamination, 4 µl of RNase was added to each tube and washing steps were performed subsequentially. DNA was eluted in 50 µl of nuclease-free water and the flow-through was reused in a second elution to increase yield. For each cockroach species, two DNA extractions were performed. All extractions were done on separate days to avoid cross-contamination between species. When necessary, DNA was cleaned by adding 50 µl of 99% ethanol and repeating the washing and elution steps.

### PCR, Sanger sequencing, and gene expression

The PCR master mix consisted of 5 µl 10x ThermoPol Reaction Buffer, 1 µl dNTP mix (10 mM/ml), 1 µl forward and 1 µl reverse primer (10 µM/ml), 0.25 µl Taq polymerase and completed with nuclease-free water to 50 µl per sample. The reaction tubes were filled with 49 µl of master mix and 1 µl of DNA. The denaturation phase was set to 30 seconds at 95°C, the annealing phase to 52°C for 60 seconds, and the elongation phase to 60 seconds at 68°C. The PCR ran for 35 cycles. PCR products were transferred to a 1.5% agar gel and run in a gel electrophoresis chamber for 35 to 45 minutes at 90V. If clear bands were visible and the negative sample was not contaminated, the samples were purified using a Qiagen PCR purification kit and sent to Eurofins sequencing facilities (https://euroainsgenomics.eu/) for Sanger sequencing. We manually curated the resulting sequences with both sequenced forward and reverse strand. Segments of sequence of low base calling confidence were replaced by the corresponding high base calling confidence segment of the reverse sequenced strand. Sequences of high quality with more than 100 bases were aligned to other *Drs* genes to validate their presence.

To quantify the expression of *Drs* genes of *D. busckii* following immune challenge, we made overnight 200rpm shaking cultures in LB, and pelleted bacteria to make pellets of *E. coli* and *M. luteus* to OD600 = 100. These pellets were then heat-killed by two cycles of 95°C (30min) and - 20°C (2h) to ensure complete bacterial killing. A dilution of each pellet was plated on LB agar to confirm bacterial killing. Pellets were then stored at -20°C until use. *Drosophila* flies were reared at 25°C with ∼50% relative humidity with a 12h light:dark cycle on the “Prop recipe” molasses food recipe [114]. Three pools of 5-7 days post-eclosion containing 5 males and 5 mated females were anaesthetized using CO2 and pricked with a 0.1mm insect pin in the thoracic pleura. Flies were allowed to recover at 25°C for 24h prior to flash freezing for later RNA extraction in TRIzol per manufacturer’s protocol.

Gene expression by qPCR was measured using the *D. melanogaster* primers and *D. busckii* primers in Data File S8. These primers specifically target the following *D. busckii* genes, given here with their orthologue in *D. melanogaster* and their LOC# identifier: *Dbus\BomBc = LOC108602262, Dbus\Dso1 = LOC108597272,* and *Dbus\RpL32 = LOC108604392.* For *Dbus\DptC,* we used primers matching 6 of the 9 *Dbus\DptC-like* loci, ignoring potential pseudogenes in designing primers. This included the following genes: *Dbus\DptC-like = LOC108595843, Dbus\DptC-like = LOC108595718, Dbus\DptC-like = LOC108597320, Dbus\DptC-like = LOC108597321, Dbus\DptC-like = LOC117134710,* and *Dbus\DptC-like = LOC117134711.* qPCR primers used 3’ ends with specificity to the target gene compared to close orthologues, and spanned intron-exon boundaries whenever possible. The 2-ΔΔCT method was used for gene expression quantification. Mastermix used was PowerUP SYBR with 5ng cDNA per 10µL reaction. qPCR was measured on a Quantstudio 3 Applied Biosystems using a fast cycle program (95°C 2min, {95°C 5s, 60°C 15s}*40, melt curve)). Expression level difference were assessed by One-way ANOVA with Tukey’s HSD.

### Sequence analysis

NF-κB binding sites in the *D. busckii Drosomycin-like* gene promoter regions were annotated using motifs for Dif (GGGHHNNDVH) and Relish (GGRDNNHHBS) from previous studies [115,116].

The sequence logo plot presented in Figure 3a was informed by a curated list of 684 Drs protein sequences recovered in our searches, removing likely pseudogenes to prevent alignment gaps. The alignment shown in Figure 3b includes just a subset of these coding sequences for visual clarity, with gaps inserted as needed to match the numbering of residues in the Figure 3a logo plot. The alignment file used to generate Figure 3a and Figure 3b can be found in Data File S9.

### Serine codon analysis

Codon alignments were performed using MAFFT plugin in Geneious Prime for a subset of *Drs* genes from across insects that were verified by manual curation to not contain potential pseudogenes. To overcome challenges of comparing species with different *Drs* gene copy number, serine codon usage for a given species used the modal codon type as representative for that residue; i.e. the most frequently occurring codon. For instance, if a species had 4 *Drs* genes and serine 3 included two TCN, one AGY, and one non-serine residue (e.g. ATA), a value of TCN was annotated for this codon. This approach included annotation of TCN-adjacent or AGY-adjacent codons in the codon table, presuming a serine ancestor. We used Bayes Factor calculations and the Jeffreys’ scale for pairwise comparisons of each lineage to assess the probability of independently deriving the same serine codon usage pattern across four codons. A log10(BF) > +2, or log10(BF) < -2 was considered as evidence of homology or convergence, respectively [73]. We required consistent patterns across most pairwise comparisons to conclude shared ancestry, as single clade-clade comparisons might skew to one trend or another through chance, while multiple comparisons skewing in the same direction was taken as support of a broader homology picture.

### Consensus sequence and percentage identity

We selected a subset of species with and without *Drs* genes to benchmark the sequence conservation of *Drs* against a widely conserved gene (Data File S10). We performed our HMM-based search on these species using an HMM profile of a partial sequence of the *dorsal* and *Dif* gene built from a sequence of similar length and intronless sequence to *Drs,* ultimately selecting the IPT_NF-κB domain. We further used the online tool EMBO-CONS (https://www.ebi.ac.uk/jdispatcher/msa/emboss_cons) to recover the consensus gene sequence for each clade. We used the Clustal-Omega online tool from the EMBO suite (https://www.ebi.ac.uk/jdispatcher/msa/clustalo) to produce a percent identity matrix using the gene sequence and the consensus sequence for each clade and calculated the average percentage identity of the genes to the consensus sequence. This gave us the average similarity of the sequences within clades. We also produced the percent identity matrix using all consensus sequences to assess the among gene sequence similarity among clades (Data File S10). Finally, we tested if the percentage identity within and among clades was higher for *Drs* genes than for the conserved genes using a CMH test using the R function *cmh_test* with the argument *alternative*=“*less”* controlling for clades and sequence length.

### Genus-level phylogeny

We built a phylogenetic tree of the species for which we screened genomes in order to test horizontal transfers of *Drs* genes in pterygote insects. We selected one species from each genus and one species per subgenus and group for the *Drosophila* genus. We selected the species with the best single copy number from the BUSCO score or the higher assembly level. We assessed the BUSCO score using BUSCO v5.3.0 with insecta_odb10 database for each of the selected species and removed any species with less than 1000 out of 1,367 BUSCO’s single copy orthologue. The set of genes which were shared by 90% of the species was concatenated using FASconCAT v1.04 [117] and aligned with MAFFT v7.505 [113]. In total the tree included 639 species and was based on 583 single copy orthologues. We reconstructed Maximum Likelihood phylogeny using IQ-TREE v 2.0.7 with substitution model of “Q.insect+G4” and default parameters. The optimal tree can be found in Data File S2.

### Gene tree reconciliation

We reconciled the gene tree with the species tree to estimate the likelihood HGT events of Drs. We first build consensus gene trees of the *Drosomycin* gene and its mature peptide based on non-redundant (i.e. identical copies were removed) *Drosomycin* genes from 65 insect species restricting the selection to one representative per genus. The amino acid sequences were first aligned using MAFFT and the resulted alignment was used to generate codon alignments from the corresponding coding nucleotide sequences using PAL2NAL v14.1 [118]. Bayesian phylogenetic analyses were performed in MrBayes v.3.2.3 using four independent analyses, including two analyses under the GTR+I+G model and two under the GTR+G model [119]. Each analysis consisted of four independent MCMC runs with 16 chains and a temperature parameter of 0.15. Analyses were run for 200 million generations, with trees and parameters sampled, and convergence assessed, every 2 million generations. The first 50 million generations were discarded as burn-in, and the remaining samples were used to construct a majority-rule consensus tree and estimate posterior probabilities. The best *Drosomycin* trees covering 95% posterior probability from each analysis were used to perform species tree gene tree reconciliation under UndatedDTL model with AleRax v1.2.0 [120,121]. The inferred HGT events were summarized across 100 reconciled gene trees, with events occurring in opposite directions between the same lineages merged into a single event. The number of times an HGT event was inferred across the 100 reconciled gene trees was used as a measure of support for that event (Data File S3). For readability, the orders of the consensus tree from each analysis were coloured using the online tree annotation tool iTOL (https://itol.embl.de/) and posterior probabilities below 75% were displayed (Figure S6).

### TE enrichment analysis

Transposable element (TE) annotations were generated for all focal genomes using EarlGrey with default parameters [122]. Analyses were conducted on 17 species representing six independent *Drs*-bearing lineages identified in this study. TE composition surrounding *Drs* loci was quantified by extracting genomic windows extending 2.5 kb upstream and downstream of each annotated gene; windows were permitted to overlap when multiple copies occurred in close proximity. TE counts were aggregated at the class level within each species. TE composition in genomic background was estimated by randomly sampling one hundred 5 kb windows from the remainder of each genome. TE enrichment in *Drs*-proximal regions was expressed as the log2 fold change relative to genomic background. The relationship between *Drs* copy number and TE abundance was evaluated across species using Pearson correlation coefficients, with P-values adjusted for multiple testing using the Benjamin-Hochberg procedure.

## Acknowledgements

This collaboration was inspired by the Royal Entomological Society conference in Penryn UK, 2023. We thank the society and conference organizers for this community event. We are grateful to Dan Hultmark for sharing his insight and *Drosomycin* sequences, including those found in Hymenoptera. The authors would like to thank the HPC Service of FUB-IT, Freie Universität Berlin and the Scientific Computation and Data Analysis Section (SCDA) of Okinawa Institute of Science and Technology, for computing time. This work was supported by German Research Foundation (DFG) grant (MC 436/5-1) to DPM, Japanese Society for the Promotion of Science (JSPS) fellowship (PE25037) to CA, and Wellcome Trust grant 227559/Z/23/Z award to MAH.

## Supplementary Material

**Supplement text**. Suspected contamination of Musca domestica samples from Zhu and Gao 2014. Mantophasmatodea transcriptome contains Drs reads suspected to come from Aromia beetle contamination. Occurrence of Drosomycin in a single representative of Tenebrionidae (Coleoptera) Supplemental text regarding the transposable elements screen of *Drosomycin* loci.

**Figure S1**. Antimicrobial peptide gene expression following infection in *D. melanogaster* and *D. busckii*.

**Figure S2**. Genomic map of the 250bp promoter region of the five *D. busckii Drosomycins* aligned to one *D. melanogaster Drs*.

**Figure S3**. Occurrence of *Drosomycin* given in the phylogenetic cladogram of the Blattodea and Mantodea orders.

**Figure S4**. Multiple sequence alignment of the mature peptide of pseudogenized *Liposcelis* sp. Drs and a selection of Drs found across other insect orders.

**Figure S5**. Outline of the required events to explain *Drs* presence across winged insects assuming a single common ancestor.

**Figure S6**. Unrooted consensus tree of the Drs sequence and mature peptide of a subset of insect species.

**Figure S7**. Transposable element (TE) composition in the genomic vicinity of *Drosomycin* (*Drs*) loci across insects.

**Figure S8**. tBLASTn sensitivity dropout given substitutions at fixed intervals.

**Table S1**. List of genomes and transcriptomes screened.

**Table S2**. Neighborhood genes of Drs in *D busckii*.

**Table S3**. Sequence found by Sanger sequencing in *Blattela germanica* and *Blatta orientalis.* **Table S4.** Correlation between transposable element classes and *Drosomycin* copy number across species.

**Data File S1.** Screenshot of the NCBI contamination warning of the two *Blattella germanica* genome assemblies.

**Data File S2.** Maximum Likelihood phylogeny of pterygote insect species in Newick format.

**Data File S3.** Overview of the inferred HGT events summarized across reconciled Drs gene trees with the species tree.

**Data File S4.** Recovered Drs sequences using HMMER approach in Fasta format for one species per genus.

**Data File S5.** Recovered Drs mature peptide sequence using HMMER approach in Fasta format for one species per genus.

**Data File S6.** Recovered Drs sequences of Collembolla species and *Lagria hirta* (Tenebrionidae).

**Data File S7.** Tabular result of Drs hits from Drosophilidae using Blast search.

**Data File S8.** List of primers used in this study and qPCR data.

**Data File S9.** Alignment of 684 curated Drs mature peptides in Fasta format.

**Data File S10.** Overview of the datasets used for testing the percentage of similarities between Drs sequences.

## Supplementary Text

### Suspected contamination of Musca domestica samples from Zhu and Gao 2014

In their 2014 paper [46], Zhu and Gao stated that two *Drosomycin* genes were found in *Musca domestica.* Since this claim, three genomes of *M. domestica* were independently sequenced. Using both HMMER and BLAST-based search methods, we did not find any hit corresponding to *Drs* in any of the three genomes. Further, the two protein and nucleotide sequences reported by Zhu and Gao are partial and lack the signal peptide, suggesting they were amplified by their PCR primer design inspired by *Drs* sequence of *D. melanogaster*. Indeed, one of the two nucleotide sequences reported by Zhu and Gao shows a 99% sequence identity with *Drosophila melanogaster* reference genome *Drs*, and the other has a 98.5% sequence identity with a *Drs-like* gene of *D. takahashii*. Taken together, we find no evidence that *M. domestica* encodes *Drs* genes, supported by three independent genome assemblies. We suspect that the sequences amplified by PCR in Zhu and Gao (2014) were contamination from *Drosophila* flies.

### Mantophasmatodea transcriptome contains *Drs* reads from *Aromia* beetle contamination

We additionally screened sequenced read archives (SRAs) from transcriptome data for lineages where genomes were unavailable. Interestingly, we recovered eight reads present in the Mantophasmatodea species *Pachyphasma brandbergense* (SRX1178919), but not other Mantophasmatodea lineages (SRX8104996, SRX1178967, SRX1178955). The fact that we recovered only eight reads suggests this could be a contamination of the *P. brandbergense* sample. We were able to assemble these reads into a complete Drs sequence. Using the assembled Drs sequence as a BLAST query, we hit on a 100% identity match to sequence from the genome of *Aromia moschata*, a longhorn beetle (Cerambycidae). We suspect a contamination of this *P. brandbergense* transcriptome explains the presence of Drosomycin in this dataset, rather than a genuine presence; this conclusion may be overturned by further sequencing, which would imply a novel HGT event into *P. brandbergense*.

We further confirm that *Drs* is present in *Aromia moschata* and other longhorn beetles, lending credibility to the hypothesis that the transcriptome SRA of *P. brandbergense* could have been contaminated by an *Aromia* beetle or related sequence.

### Occurrence of *Drosomycin* in a single representative of Tenebrionidae (Coleoptera)

During the extended screen of Coleoptera genomes, we recovered *Drs* hits in *Lagria hirta* (Tenebrionidae). The sequences recovered come from two assemblies GCA_947359435 and GCA_947359425 derived from the same BioSample SAMEA11025111. One of the assemblies is of chromosome level with BUSCO completeness of 98.5% and the sequences recovered were not seemingly from a contamination of another known insect (e.g. *Drosophila* or beetles). Yet, since none of the 12 other Tenebrionidae genomes available have *Drs* genes and the two assemblies are not independent from each other, we prefer to present this finding conservatively, and so do not include this sequence as a confirmed *Drs* presence in this lineage. However, we provide the sequence in the supplementary (Data File S6). As more genomes of *Lagria* and relatives become available, they may strengthen this finding, or possibly suggest it stems from a contamination of the *Lagria* genome sequence.

### Transposable Elements screen of *Drosomycin* loci

The repeated horizontal acquisition of the *Drs* locus across insect orders raises the question of whether a shared transposable element (TE) mediated mechanism could underlie its insertion into diverse host genomes. To address this, we examined patterns of TE composition in 5 kb windows flanking *Drs* loci across 17 species representing six putatively independent Drs-bearing lineages identified in this study. If *Drs* insertions are derived from a common source population or a conserved molecular mechanism, we might expect to observe a consistent enrichment of particular TE families or classes in the vicinity of *Drs* loci across species.

Across species, simple repeats were detected within 5 kb of *Drs* loci in all examined genomes. In addition, satellite repeats, DNA/TcMar-Mariner elements, and LINE/RTE-BovB elements were found in proximity to *Drs* loci in a majority of species (at least 10 of 17; Figure S7). To assess whether these observations reflected a specific association with *Drs* loci rather than general properties of host genomes, we compared TE abundance in *Drs*-proximal windows to randomly sampled genomic windows of equivalent size within each species. This analysis revealed no significant enrichment of any TE family or class in the vicinity of *Drs* loci relative to genome-wide background levels. Thus, while multiple TE classes frequently co-occur with *Drs* loci, their local abundance does not differ from expectation given overall genomic TE composition.

We further examined the relationship between *Drs* copy number and local TE content across species, finding some positive correlations between LINE/I-Jockey and DNA/TcMar-Tc1 elements in specific cases (Table S4; Figure S7). However, when TE abundance in *Drs*-proximal windows was compared to randomly sampled genomic windows of equivalent size, none of these TE classes were consistently enriched near *Drs* loci across species (Figure S7).

Taken together, our TE screen provides no evidence for a shared transposable element or source population responsible for the horizontal transfer of *Drs* across insect lineages. In most cases, TE composition near to *Drs* loci in a given species was comparable to TE composition in the rest of the genome. Given the deep evolutionary timescales involved and the rapid turnover of transposable elements, any TE signatures that might have been associated with insertion events may have been eroded beyond recognition.

---

**Figure S1.**
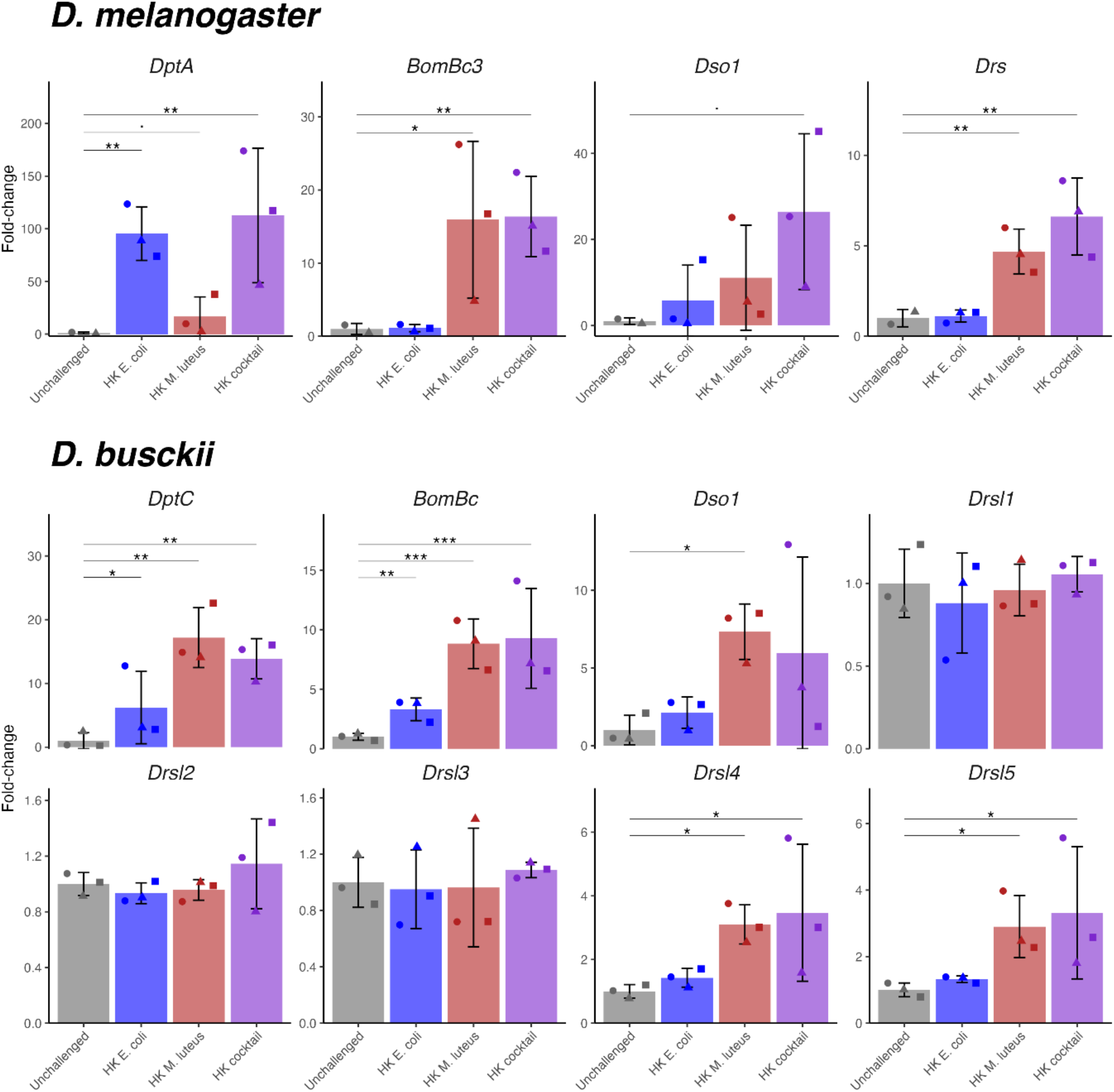
Antimicrobial gene expression following infection in *Drosophila melanogaster* and *D. busckii*. The *D. busckii* 2R *DbDrsl4* and *DbDrsl5* are immune-induced by heat-killed M. luteus injection similar to Toll-regulated Bomanins (BomBc) and *D. melanogaster Drs*. In contrast, none of the 2L *DbDrs1-3* were immune induced by the pathogen tested.

**Figure S2.**
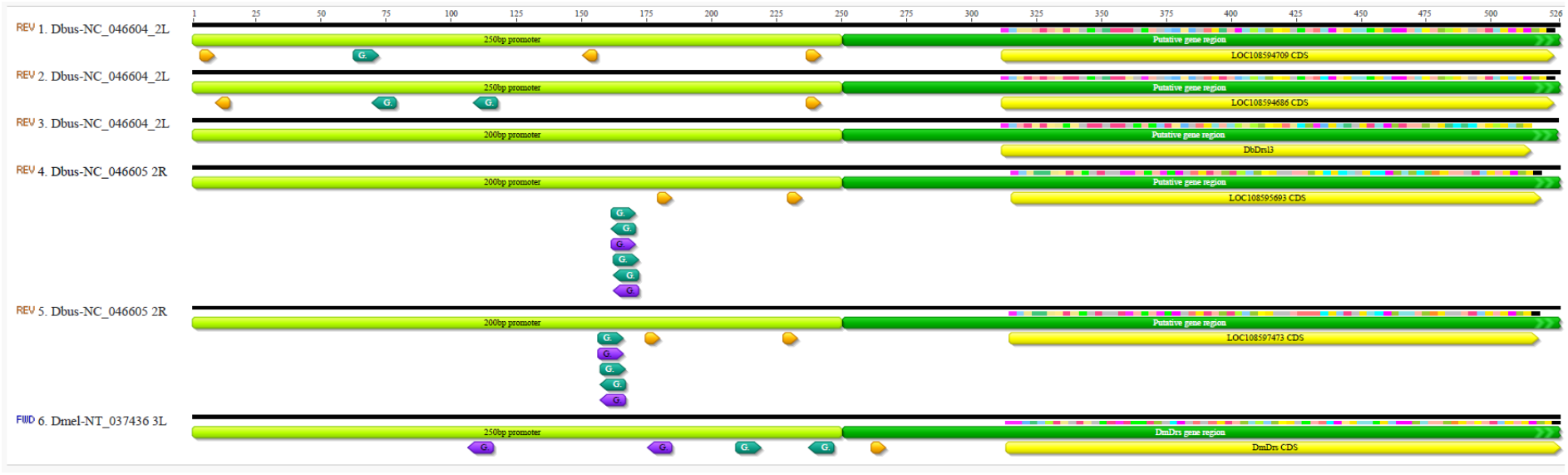
Genomic map of the 250bp promoter region of the Uive *D. busckii Drosomycins* aligned to *D. melanogaster Drs*. Binding sites (purple arrows) for the NF-κB protein Dorsal-related immunity factor (Dif) are present in the promoter region (light green) of the putative Drs region (dark green) of the 2R *DbDrsl4* and *DbDrsl5,* as found in *D. melanogaster Drs*, but not in the 2L *DbDrsl1*, *DbDrsl2*, or *DbDrsl3.* The coding sequence (CDS) of each *Drs* is highlighted in yellow. Turquoise arrows correspond to Rel-matching sites and yellow-orange arrows correspond to WGATAR-matching GATA sites [123].

**Figure S3:**
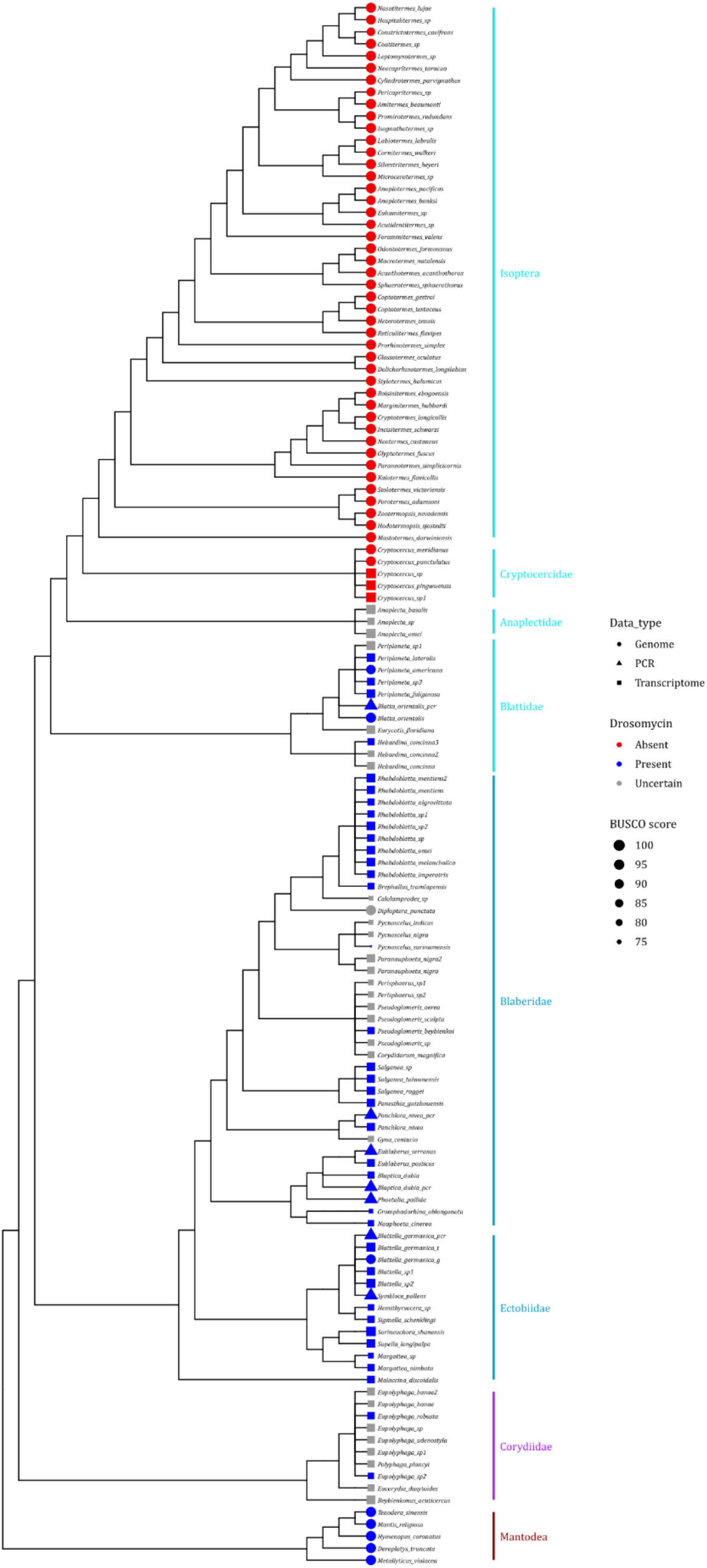
Occurrence of Drosomycin in the phylogeny of Blattodea and Mantodea. The size of the branch tip is proportional to the BUSCO completeness score. Blue tips indicate that Drosomycin-like gene(s) were found, red tips indicate that there is credible evidence for their absence (i.e. no hit found in high resolution genome assemblies), grey tips indicate that the gene was not found but the support for true absence is low. Drosomycin-like gene(s) were screened in genomic (circle tip), or transcriptomic (square tips) data or confirmed by PCR and Sanger sequencing (triangle tips). The abundance of positive hits in most Mantodea and Blattodea clades suggests that Drosomycin was gained in the common ancestor and lost secondarily in Isoptera and Cryptocercidae. Cladogram drawn from [63–65].

**Figure S4.**
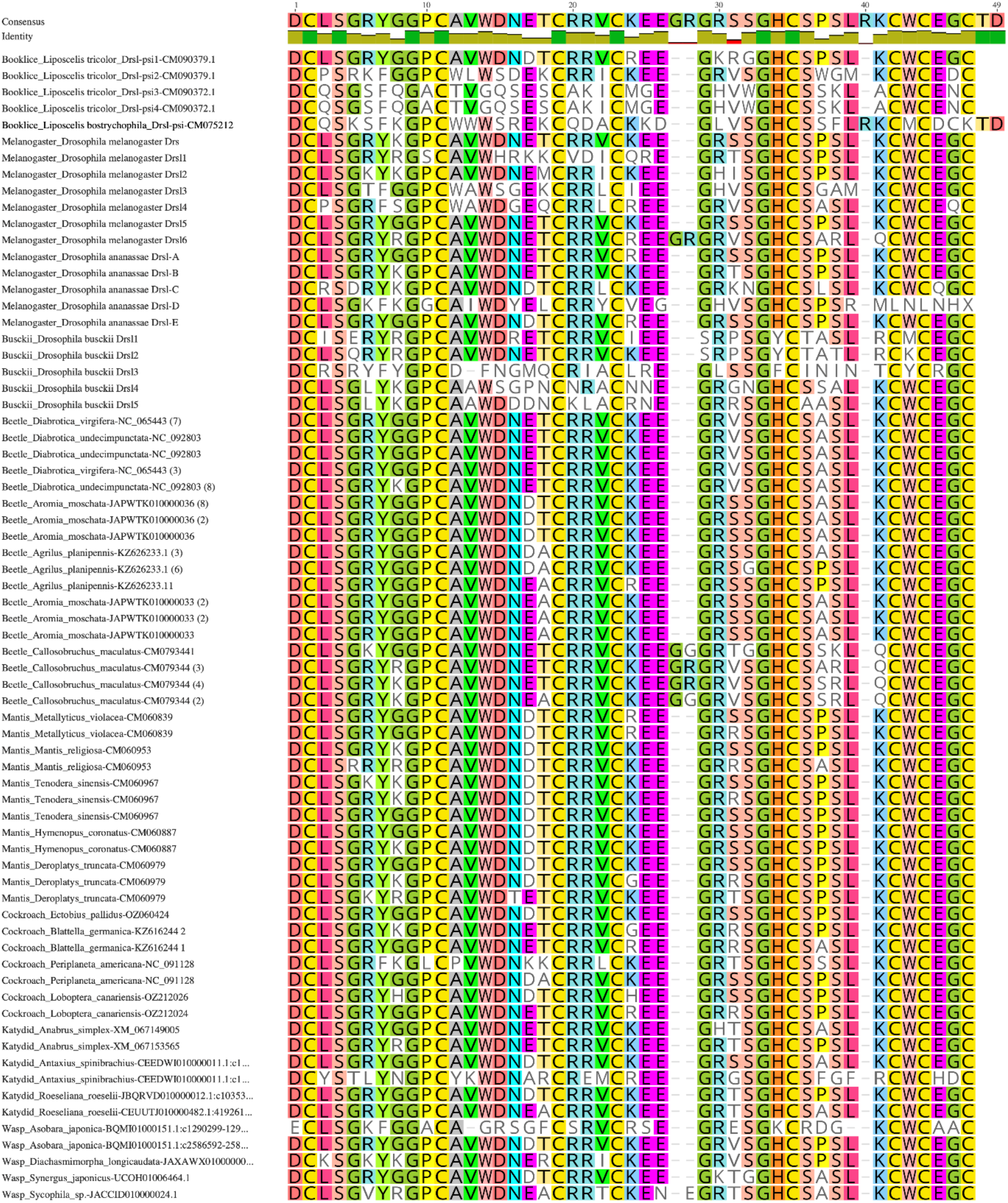
Multiple sequence alignment of the mature peptide of pseudogenized *Liposcelis* sp. Drs and a selection of Drs found across other insect orders. The Liposcelis Drs (top of the panel) present all the Cystein codon to be defined as Drs-like peptide. However, the sequences are clearly degraded and are missing one or two of the expected serine codons.

**Figure S5.**
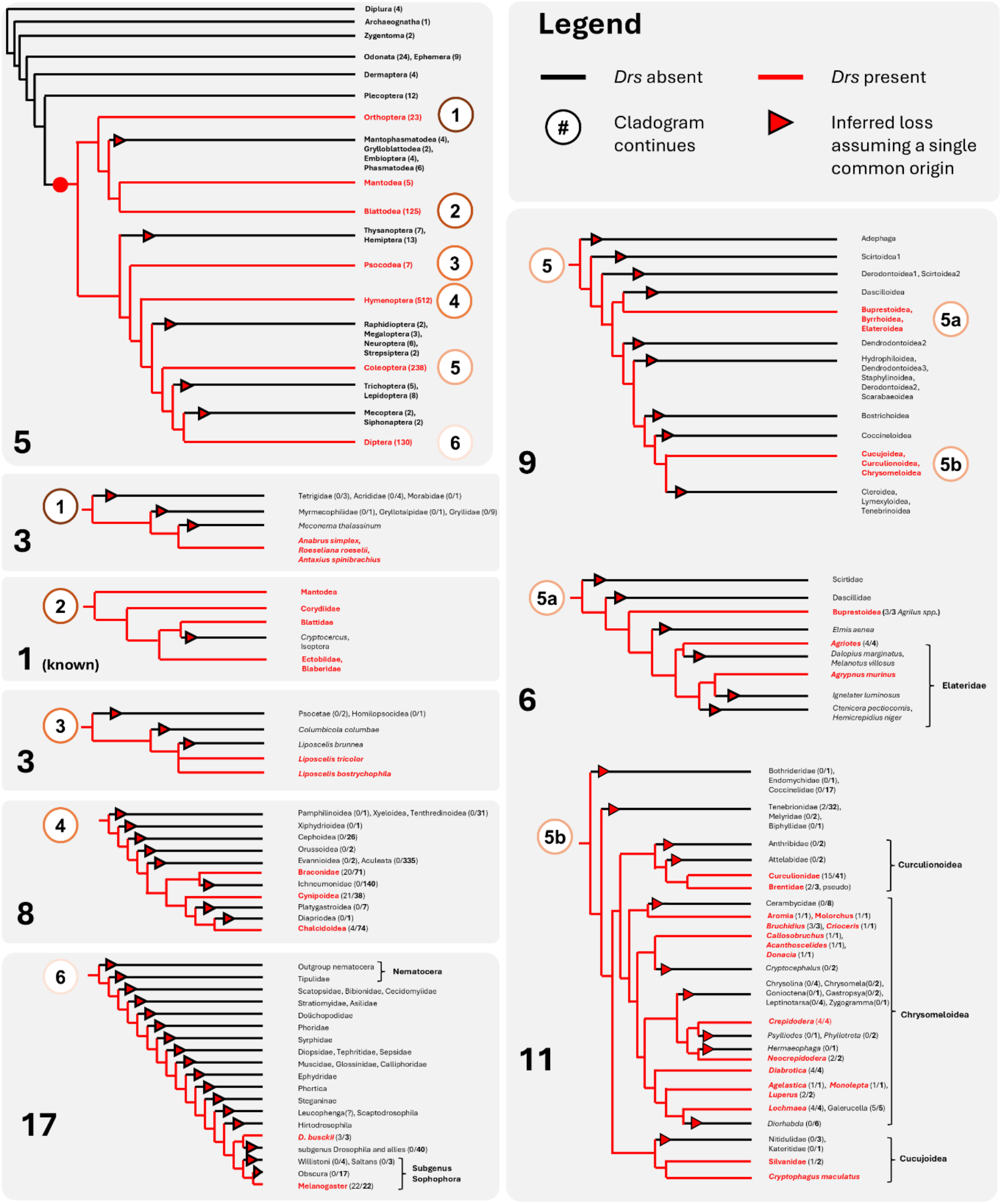
Outline of the required events to explain *Drs* presence across winged insects assuming a single common ancestor. Imagining the Drs genes of all insects occur through inheritance from a single common ancestral sequence requires 1 gain (red circle) and 63 independent loss events (triangles) per the current understanding of insect phylogenetics: this includes five losses across major insect orders (top left panel), three losses within Orthoptera, one loss in Blattodea (a genuine secondary loss), three losses in Psocodea, 26 losses within Coleoptera (some of which are likely genuine secondary losses), 8 losses within Hymenoptera, and 17 losses within Diptera. In the top left panel, the Figure in brackets corresponds to the number of species screened for a given order. In the other panels, the ratio in brackets corresponds to the number of assemblies with *Drs* genes by the total number of assemblies screened for the clade. Modern phylogenies were used for the cladogram of Insecta [67], Orthoptera [124], Blattodea [63,64], Psocodea [125], Hymenoptera [126], Coleoptera [68,127,128], Diptera [47,51] and Drosophilidae [48].

**Figure S6.**
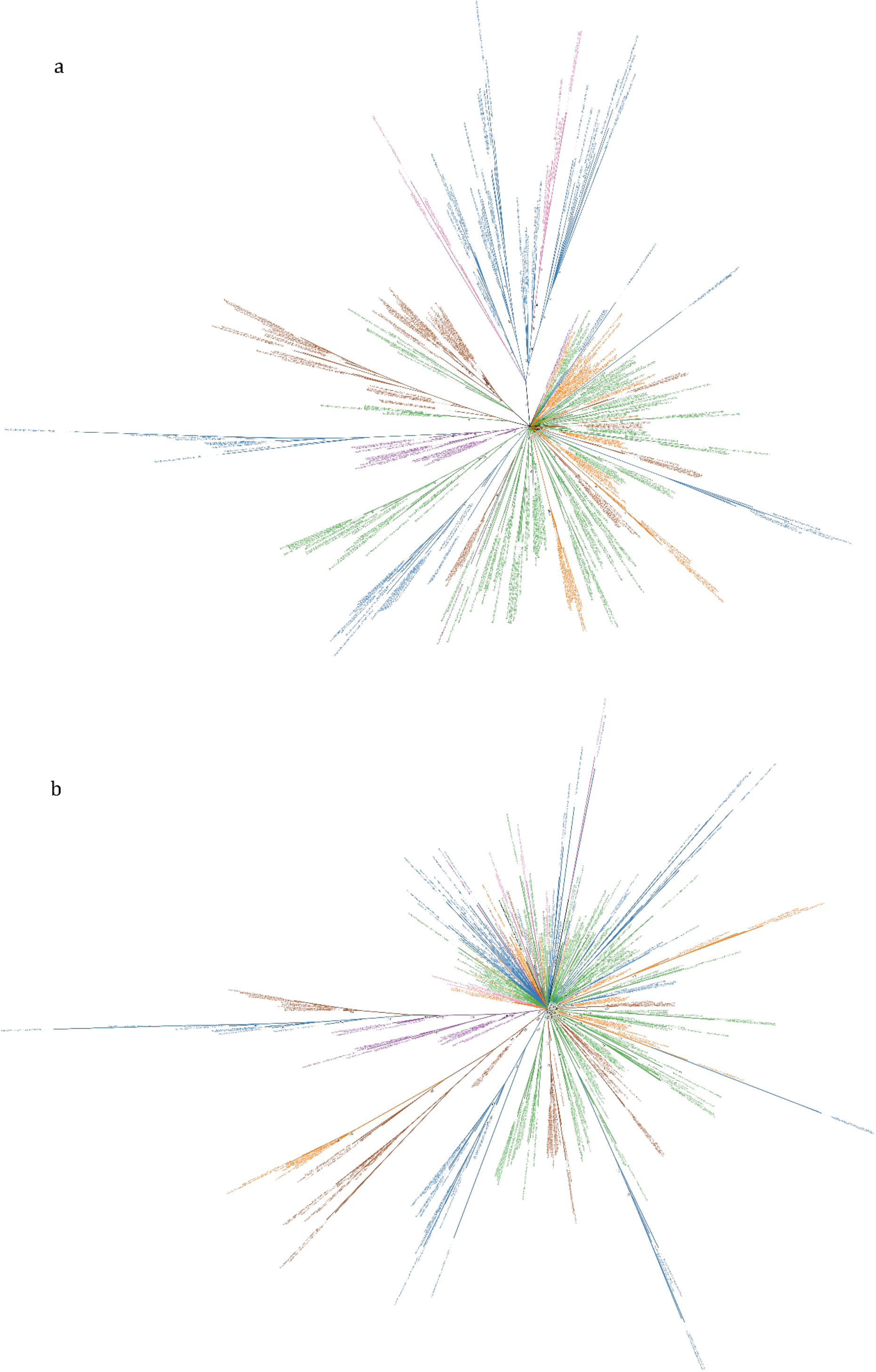
Unrooted consensus tree of (a) the Drs sequence and (b) mature peptide of a subset of insect species. The tree was produced with MrBayes v3.2.3 using codon alignment of the Drs sequences from one species per genus analyzed in this study (see Method). Posterior probabilities below 75% are displayed on branch labels. Several paraphyly across orders are observable on the full sequence Drs tree, noteworthily, between wasps (blue) and flies (pink), beetles (green) and cockroaches (brown), and katydids (purple) and wasps. The mature peptide tree is much less informative presenting only a few paraphyly, each involving the wasps.

**Figure S7.**
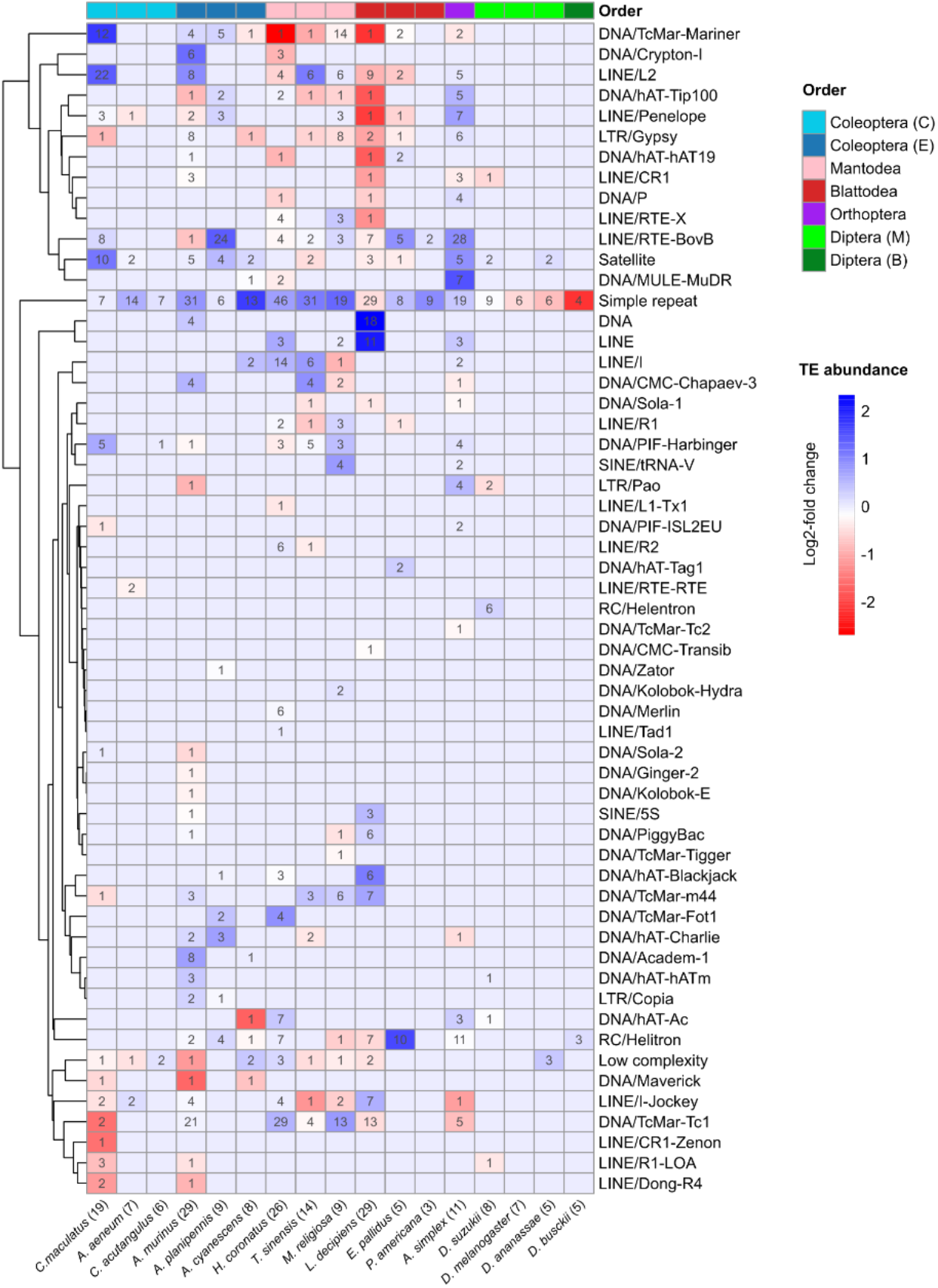
Transposable element (TE) composition in the genomic vicinity of *Drosomycin* (*Drs*) loci across insects. Heatmap showing the log2-fold change (log2FC) in abundance of TE classes within 5 kb windows flanking *Drs* loci, relative to randomly sampled 5 kb windows from the remainder of each genome. Each cell represents a species–TE class combination. Cell colour indicates log2FC values, while numbers within cells denote the total number of TE copies of the corresponding class detected within *Drs*-proximal windows for that species. The number of *Drs* gene copies identified in each genome is shown in parentheses following the species name. Species are grouped according to the six insect lineages represented in this study. Within Coleoptera, Coleoptera (C) denotes species from the Cucujiformia infraorder, while Coleoptera (E) denotes species from the Elateriformia infraorder. Within Diptera, Diptera (M) corresponds to species from the *melanogaster* species group, and Diptera (B) corresponds to *Drosophila busckii*. TE annotations for all genomes were generated using EarlGrey [122].

**Figure S8.**
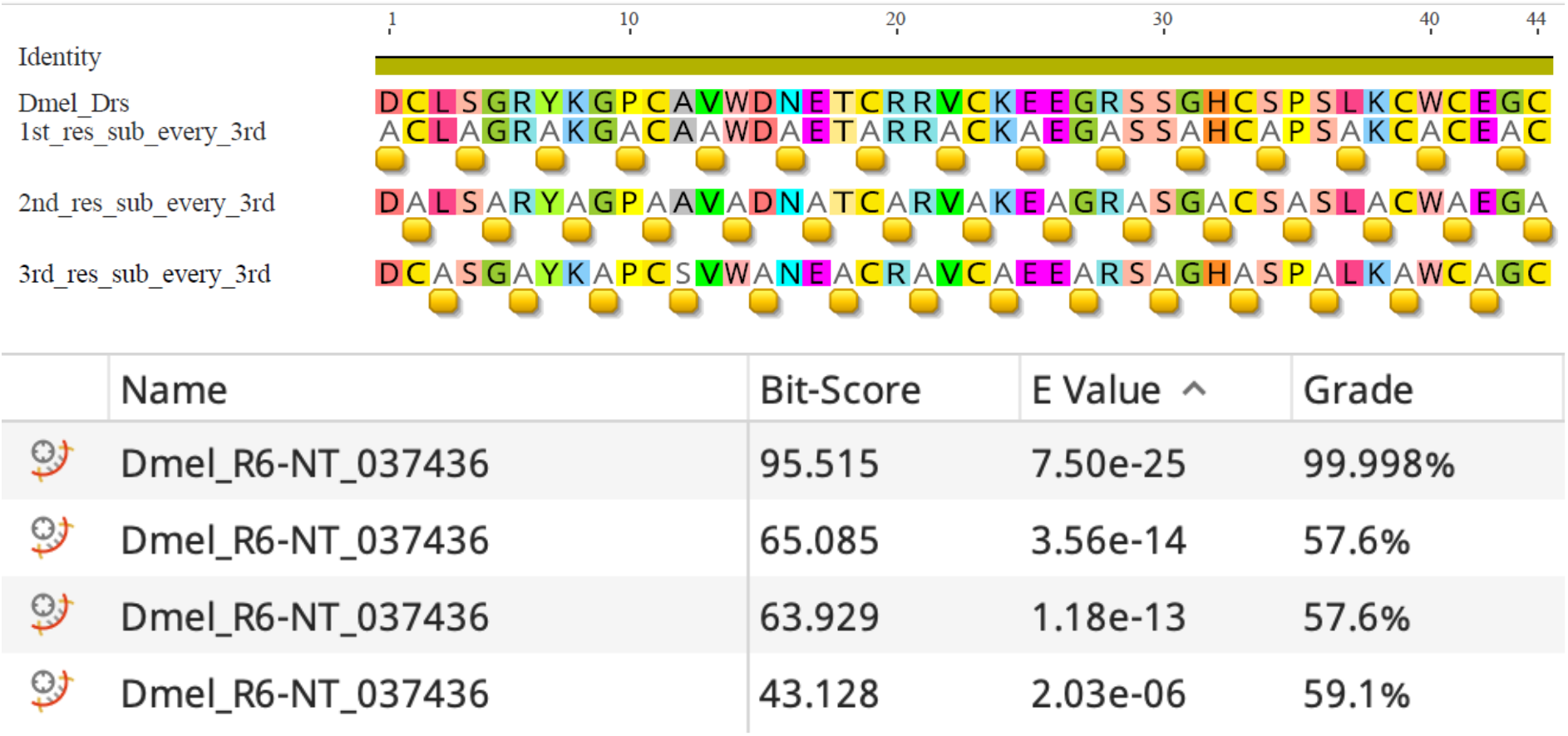
tBLASTn sensitivity dropout given substitutions at Uixed intervals. The E-Value of the hit decays rapidly depending on the number and position of alanine every three residues. Furthermore, due to short sequence, even an unadulterated Dmel\Drs query returns just 7.5e-25, and so artificially high E-values and manual curation are essential in querying for short AMP sequences across species.

## Notes

### Competing Interest Statement

The authors have declared no competing interest.

### Summary of Updates

Minor revision to address incongruent number of insect orders reported

## Literature Cited

1. One Health High-Level Expert Panel, Adisasmito WB, Almuhairi S, Behravesh CB, Bilivogui P, Bukachi SA, et al. One Health: A new definition for a sustainable and healthy future. PLoS Pathog. 2022;18: e1010537. doi:10.1371/journal.ppat.1010537

2. Shultz AJ, Sackton TB. Immune genes are hotspots of shared positive selection across birds and mammals. eLife. 2019;8: e41815. doi:10.7554/eLife.41815

3. Clark AG, Eisen MB, Smith DR, Bergman CM, Oliver B, Markow TA, et al. Evolution of genes and genomes on the *Drosophila* phylogeny. Nature. 2007;450: 203–218. doi:10.1038/nature06341

4. Jones JDG, Dangl JL. The plant immune system. Nature. 2006;444: 323–329. doi:10.1038/nature05286

5. Klimovich A, Bosch TCG. Novel technologies uncover novel ‘anti’-microbial peptides in *Hydra* shaping the species-specific microbiome. Phil Trans R Soc B. 2024;379: 20230058. doi:10.1098/rstb.2023.0058

6. Pradeu T, Thomma BPHJ, Girardin SE, Lemaitre B. The conceptual foundations of innate immunity: Taking stock 30 years later. Immunity. 2024;57: 613–631. doi:10.1016/j.immuni.2024.03.007

7. Balaji SK, Balasundarasekar B, Khuwaja WM, Dolan KM, Dong X. Antimicrobial peptide signaling in skin diseases. JID Innov. 2025;5: 100354. doi:10.1016/j.xjidi.2025.100354

8. Hanson MA, Westlake HE, Bulet P, Lemaitre B. The antimicrobial and host defense peptides of *Drosophila melanogaster*. Annu Rev Microbiol. 2025;79: 335–360. doi:10.1146/annurev-micro-101923-100221

9. Unckless RL, Howick VM, Lazzaro BP. Convergent balancing selection on an antimicrobial peptide in *Drosophila*. Curr Biol. 2016;26: 257–262. doi:10.1016/j.cub.2015.11.063

10. Myers AN, Lawhon SD, Diesel AB, Bradley CW, Rodrigues Hoffmann A, Murphy WJ, et al. An ancient haplotype containing antimicrobial peptide gene variants is associated with severe fungal skin disease in Persian cats. PLoS Genet. 2022;18: e1010062. doi:10.1371/journal.pgen.1010062

11. Hanson MA. When the microbiome shapes the host: immune evolution implications for infectious disease. Phil Trans R Soc B. 2024;379: 20230061. doi:10.1098/rstb.2023.0061

12. Imler J-L, Cai H, Meignin C, Martins N. Evolutionary immunology to explore original antiviral strategies. Philos Trans R Soc Lond B Biol Sci. 2024;379: 20230068. doi:10.1098/rstb.2023.0068

13. Kaessmann H. Origins, evolution, and phenotypic impact of new genes. Genome Res. 2010;20: 1313–1326. doi:10.1101/gr.101386.109

14. Moran NA, Jarvik T. Lateral transfer of genes from fungi underlies carotenoid production in aphids. Science. 2010;328: 624–627. doi:10.1126/science.1187113

15. Dudzic JP, Hanson MA, Iatsenko I, Kondo S, Lemaitre B. More than black or white: Melanization and Toll share regulatory serine proteases in *Drosophila*. Cell Rep. 2019;27: 1050–1061.e3. doi:10.1016/j.celrep.2019.03.101

16. Hanson MA, Grollmus L, Lemaitre B. Ecology-relevant bacteria drive the evolution of host antimicrobial peptides in *Drosophila*. Science. 2023;381: eadg5725. doi:10.1126/science.adg5725

17. Hedengren M, Borge K, Hultmark D. Expression and evolution of the *Drosophila* attacin/diptericin gene family. Biochem Biophys Res Commun. 2000;279: 574–81. doi:10.1006/bbrc.2000.3988

18. Hanson MA, Hedelin L. Humoral immunity in insects: Antimicrobial peptides and other host defense peptides. Reference Module in Life Sciences. Elsevier; 2025. p. B9780323954242000029. doi:10.1016/B978-0-323-95424-2.00002-9

19. Casteels P, Ampe C, Riviere L, Damme J, Elicone C, Fleming M, et al. Isolation and characterization of abaecin, a major antibacterial response peptide in the honeybee (*Apis mellifera*). Eur J Biochem. 1990;187: 381–386. doi:10.1111/j.1432-1033.1990.tb15315.x

20. Cociancich S, Dupont A, Hegy G, Lanot R, Holder F, Hetru C, et al. Novel inducible antibacterial peptides from a hemipteran insect, the sap-sucking bug *Pyrrhocoris apterus*. Biochem J. 1994;300: 567–575. doi:10.1042/bj3000567

21. Bulet P, Urge L, Ohresser S, Hetru C, Otvos L. Enlarged scale chemical synthesis and range of activity of drosocin, an O-glycosylated antibacterial peptide of *Drosophila*. Eur J Biochem. 1996;238: 64–69. doi:10.1111/j.1432-1033.1996.0064q.x

22. Mardirossian M, Pérébaskine N, Benincasa M, Gambato S, Hofmann S, Huter P, et al. The dolphin proline-rich antimicrobial peptide Tur1A inhibits protein synthesis by targeting the bacterial ribosome. Cell Chem Biol. 2018;25: 530–539.e7. doi:10.1016/j.chembiol.2018.02.004

23. Lachat J, Lextrait G, Jouan R, Boukherissa A, Yokota A, Jang S, et al. Hundreds of antimicrobial peptides create a selective barrier for insect gut symbionts. Proc Natl Acad Sci USA. 2024;121: e2401802121. doi:10.1073/pnas.2401802121

24. Conlon JM, Kolodziejek J, Nowotny N. Antimicrobial peptides from ranid frogs: taxonomic and phylogenetic markers and a potential source of new therapeutic agents. Biochim Biophys Acta - Proteins Proteom. 2004;1696: 1–14. doi:10.1016/j.bbapap.2003.09.004

25. Lima AM, Azevedo MIG, Sousa LM, Oliveira NS, Andrade CR, Freitas CDT, et al. Plant antimicrobial peptides: An overview about classification, toxicity and clinical applications. Int J Biol Macromol. 2022;214: 10–21. doi:10.1016/j.ijbiomac.2022.06.043

26. Augustin R, Schröder K, Murillo Rincón AP, Fraune S, Anton-Erxleben F, Herbst E-M, et al. A secreted antibacterial neuropeptide shapes the microbiome of *Hydra*. Nat Commun. 2017;8: 698. doi:10.1038/s41467-017-00625-1

27. Marra A, Hanson MA, Kondo S, Erkosar B, Lemaitre B. *Drosophila* antimicrobial peptides and lysozymes regulate gut microbiota composition and abundance. mBio. 2021; e0082421. doi:10.1128/mBio.00824-21

28. Broekaert WF, Terras FR, Cammue BP, Osborn RW. Plant defensins: novel antimicrobial peptides as components of the host defense system. Plant physiol. 1995;108: 1353–1958.

29. Gerdol M, Schmitt P, Venier P, Rocha G, Rosa RD, Destoumieux-Garzón D. Functional insights from the evolutionary diversification of Big Defensins. Front Immunol. 2020;11: 758. doi:10.3389/fimmu.2020.00758

30. Shafee TMA, Lay FT, Phan TK, Anderson MA, Hulett MD. Convergent evolution of defensin sequence, structure and function. Cell Mol Life Sci. 2017;74: 663–682. doi:10.1007/s00018-016-2344-5

31. Hultmark D, Steiner H, Rasmuson T, Boman HG. Insect Immunity. Purification and properties of three inducible bactericidal proteins from hemolymph of immunized pupae of *Hyalophora cecropia*. Eur J Biochem. 1980;106: 7–16. doi:10.1111/j.1432-1033.1980.tb05991.x

32. Quesada H, Ramos-Onsins SE, Aguade M. Birth-and-death evolution of the Cecropin multigene family in *Drosophila*. J Mol Evol. 2005;60: 1–11. doi:10.1007/s00239-004-0053-4

33. Soucy SM, Huang J, Gogarten JP. Horizontal gene transfer: building the web of life. Nat Rev Genet. 2015;16: 472–482. doi:10.1038/nrg3962

34. Houck MA, Clark JB, Peterson KR, Kidwell MG. Possible Horizontal Transfer of *Drosophila* Genes by the Mite *Proctolaelaps regalis*. Science. 1991;253: 1125–1128. doi:10.1126/science.1653453

35. Nakabachi A. Horizontal gene transfers in insects. Curr Opi Insect Sci. 2015;7: 24–29. doi:10.1016/j.cois.2015.03.006

36. Shelomi M, Danchin EGJ, Heckel D, Wipfler B, Bradler S, Zhou X, et al. Horizontal gene transfer of pectinases from bacteria preceded the diversification of stick and leaf insects. Sci Rep. 2016;6: 26388. doi:10.1038/srep26388

37. Liu C, Hellemans S, Kinjo Y, Mikhailova AA, Aumont C, Weng Y-M, et al. Recurrent horizontal gene transfers across diverse termite genomes. Evolution. 2026; qpag003. doi:10.1093/evolut/qpag003

38. Verster KI, Wisecaver JH, Karageorgi M, Duncan RP, Gloss AD, Armstrong EE, et al. Horizontal transfer of bacterial cytolethal distending toxin B genes to insects. Mol Biol Evol. 2019;36: 2105–2110. doi:10.1093/molbev/msz146

39. La Camera S, Gouzerh G, Dhondt S, Hoffmann L, Fritig B, Legrand M, et al. Metabolic reprogramming in plant innate immunity: The contributions of phenylpropanoid and oxylipin pathways. Immunol Rev. 2004;198: 267–284. doi:10.1111/j.0105-2896.2004.0129.x

40. Perlmutter JI, Chapman JR, Wilkinson MC, Nevarez-Saenz I, Unckless RL. A single amino acid polymorphism in natural Metchnikowin alleles of *Drosophila* results in systemic immunity and life history tradeoffs. PLoS Genet. 2024;20: e1011155. doi:10.1371/journal.pgen.1011155

41. Rommelaere S, Carboni A, Bada Juarez JF, Boquete J-P, Abriata LA, Meireles FTP, et al. A humoral stress response protects *Drosophila* tissues from antimicrobial peptides. Curr Biol. 2024; 1–12. doi:10.1101/2023.07.24.550293

42. Bacalum M, Radu M. Cationic antimicrobial peptides cytotoxicity on mammalian cells: An analysis using therapeutic index integrative concept. Int J Pept Res Ther. 2015;21: 47–55. doi:10.1007/s10989-014-9430-z

43. Fehlbaum P, Bulet P, Michaut L, Lagueux M, Broekaert WF, Hetru C, et al. Insect immunity: Septic injury of *Drosophila* induces the synthesis of a potent antifungal peptide with sequence homology to plant antifungal peptides. J Biol Chem. 1994;269: 33159–33163. doi:10.1016/S0021-9258(20)30111-3

44. Jiggins FM. The evolution of antifungal peptides in *Drosophila*. Genetics. 2005;171: 1847– 1859. doi:10.1534/genetics.105.045435

45. Cammue BP, De Bolle MF, Terras FR, Proost P, Van Damme J, Rees SB, et al. Isolation and characterization of a novel class of plant antimicrobial peptides form *Mirabilis jalapa* L. seeds. J Biol Chem. 1992;267: 2228–2233. doi:10.1016/S0021-9258(18)45866-8

46. Zhu S, Gao B. Nematode-derived drosomycin-type antifungal peptides provide evidence for plant-to-ecdysozoan horizontal transfer of a disease resistance gene. Nat Commun. 2014;5: 3154. doi:10.1038/ncomms4154

47. Hanson MA, Lemaitre B, Unckless RL. Dynamic evolution of antimicrobial peptides underscores trade-offs between immunity and ecological fitness. Front Immunol. 2019;10: 2620. doi:10.3389/fimmu.2019.02620

48. Kim BY, Gellert HR, Church SH, Suvorov A, Anderson SS, Barmina O, et al. Single-aly genome assemblies fill major phylogenomic gaps across the Drosophilidae Tree of Life. PLoS Biol. 2024;22: e3002697. doi:10.1371/journal.pbio.3002697

49. Suvorov A, Kim BY, Wang J, Armstrong EE, Peede D, D’Agostino ERR, et al. Widespread introgression across a phylogeny of 155 *Drosophila* genomes. Curr Biol. 2021; S0960982221014962. doi:10.1016/j.cub.2021.10.052

50. Thiébaut A, Altenhoff AM, Campli G, Glover N, Dessimoz C, Waterhouse RM. DrosOMA: the *Drosophila* Orthologous Matrix browser. F1000Res. 2024;12: 936. doi:10.12688/f1000research.135250.2

51. Vicoso B, Bachtrog D. Numerous transitions of sex chromosomes in Diptera. PLoS Biol. 2015;13. doi:10.1371/journal.pbio.1002078

52. He S, Sieksmeyer T, Che Y, Mora MAE, Stiblik P, Banasiak R, et al. Evidence for reduced immune gene diversity and activity during the evolution of termites. Proc R Soc B: Biol Sci. 2021;288: 20203168. doi:10.1098/rspb.2020.3168

53. Liu C, Aumont C, Mikhailova AA, Audisio T, Hellemans S, Weng Y-M, et al. Unravelling the evolution of wood-feeding in termites with 47 high-resolution genome assemblies. Nat Commun. 2025;16: 11154. doi:10.1038/s41467-025-65969-5

54. Harrison MC, Arning N, Kremer LPM, Ylla G, Belles X, Bornberg-Bauer E, et al. Expansions of key protein families in the German cockroach highlight the molecular basis of its remarkable success as a global indoor pest. J Exp Zool B: Mol Dev Evol. 2018;330: 254–264. doi:10.1002/jez.b.22824

55. Silva FJ, Muñoz-Benavent M, García-Ferris C, Latorre A. *Blattella germanica* displays a large arsenal of antimicrobial peptide genes. Sci Rep. 2020;10: 21058. doi:10.1038/s41598-020-77982-3

56. Jones AR, Mikhailova AA, Aumont C, Berger J, Liu C, He S, et al. *Cryptocercus* genomes expand knowledge of adaptations to xylophagy and termite sociality. Genome Biol Evol. 2026; evag028. doi:10.1093/gbe/evag028

57. Liu J, Zhang J, Han W, Wang Y, He S, Wang Z. Advances in the understanding of Blattodea evolution: Insights from phylotranscriptomics and spermathecae. Mol Phylogenetics Evol. 2023;182: 107753. doi:10.1016/j.ympev.2023.107753

58. Fouks B, Harrison MC, Mikhailova AA, Marchal E, English S, Carruthers M, et al. Live-bearing cockroach genome reveals convergent evolutionary mechanisms linked to viviparity in insects and beyond. Iscience. 2023;26. doi:10.1016/j.isci.2023.107832

59. Dockman RL, Simmonds TJ, Vogel KJ, Geib SM, Ottesen EA. Improved chromosome-level genome assembly of the American cockroach, *Periplaneta americana*. G3: Genes Genomes Genet. 2026;16: jkaf247. doi:10.1093/g3journal/jkaf247

60. Hunter T, Natural History Museum Genome Acquisition Lab, Wellcome Sanger Institute Tree Life, Darwin Tree of Life Consortium. The genome sequence of the tawny cockroach, *Ectobius (Ectobius) pallidus* (Olivier, 1789). Wellcome Open Res. 2025;10: 22. doi:10.12688/wellcomeopenres.23463.1

61. López H, Oromí P, Belles X, Ylla G, Escudero N, Fernández R, et al. ERGA-BGE Reference Genome of *Loboptera subterranea*: A cave adapted cockroach endemic to the Canary Islands. bioRxiv; 2025. doi:10.1101/2025.02.06.636782

62. Liu H, Lei L, Jiang F, Zhang B, Wang H, Zhang Y, et al. The genomes of 5 mantises provide insights into sex chromosome evolution and Mantodea phylogeny clarification. GigaScience. 2026;15: giaf158. doi:10.1093/gigascience/giaf158

63. Wang Z, Shi Y, Qiu Z, Che Y, Lo N. Reconstructing the phylogeny of Blattodea: robust support for interfamilial relationships and major clades. Sci Rep. 2017;7: 3903. doi:10.1038/s41598-017-04243-1

64. Djernæs M, Murienne J. Phylogeny of Blattoidea (Dictyoptera: Blattodea) with a revised classification of Blattidae. Arthropod Syst Phylogeny. 2022;80: 209–228. doi:10.3897/asp.80.e75819

65. Ma Y, Zhang L-P, Lin Y-J, Yu D-N, Storey KB, Zhang J-Y. Phylogenetic relationships and divergence dating of Mantodea using mitochondrial phylogenomics. Syst Entomol. 2023;48: 644–657. doi:10.1111/syen.12596

66. Eddy SR. Accelerated profile HMM searches. PLoS Comput Biol. 2011;7: e1002195. doi:10.1371/journal.pcbi.1002195

67. Misof B, Liu S, Meusemann K, Peters RS, Donath A, Mayer C, et al. Phylogenomics resolves the timing and pattern of insect evolution. Science. 2014;346: 763–767. doi:10.1126/science.1257570

68. Zhang S-Q, Che L-H, Li Y, Dan Liang, Pang H, Sâlipiński A, et al. Evolutionary history of Coleoptera revealed by extensive sampling of genes and species. Nat Commun. 2018;9:205. doi:10.1038/s41467-017-02644-4

69. Li Y-D, Xie Z-Y, Du Y-L, Zhou Z, Mao X-M, Lv L-X, et al. The rapid evolution of signal peptides is mainly caused by relaxed selection on non-synonymous and synonymous sites. Gene. 2009;436: 8–11. doi:10.1016/j.gene.2009.01.015

70. Kumar S, Stecher G, Suleski M, Hedges SB. TimeTree: A resource for timelines, timetrees, and divergence times. Mol Biol Evol. 2017;34: 1812–1819. doi:10.1093/molbev/msx116

71. Lemaitre B, Nicolas E, Michaut L, Reichhart J-M, Hoffmann JA. The dorsoventral regulatory gene cassette spätzle/Toll/cactus controls the potent antifungal response in *Drosophila* adults. Cell. 1996;86: 973–983. doi:10.1016/S0092-8674(00)80172-5

72. Rogozin IB, Belinky F, Pavlenko V, Shabalina SA, Kristensen DM, Koonin EV. Evolutionary switches between two serine codon sets are driven by selection. Proc Natl Acad Sci USA. 2016;113: 13109–13113. doi:10.1073/pnas.1615832113

73. Jeffreys H. The theory of probability. 3rd ed. Oxford: Oxford University Press; 1961. doi:10.1093/oso/9780198503682.001.0001

74. Hanson MA, Hamilton PT, Perlman SJ. Immune genes and divergent antimicrobial peptides in flies of the subgenus *Drosophila*. BMC Evol Biol. 2016;16: 228. doi:10.1186/s12862-016-0805-y

75. Hollox EJ, Armour JAL. Directional and balancing selection in human beta-defensins. BMC Evol Biol. 2008;8. doi:10.1186/1471-2148-8-113

76. Arunkumar R, Zhou SO, Day JP, Bakare S, Pitton S, Zhang Y, et al. Natural selection has driven the recurrent loss of an immunity gene that protects *Drosophila* against a major natural parasite. Proc Natl Acad Sci USA. 2023;120: e2211019120. doi:10.1073/pnas.2211019120

77. Hanson MA, Kondo S, Lemaitre B. *Drosophila* immunity: the Drosocin gene encodes two host defence peptides with pathogen-specific roles. Proc Biol Sci. 2022;289: 20220773. doi:10.1098/rspb.2022.0773

78. Simon A, Kullberg BJ, Tripet B, Boerman OC, Zeeuwen P, Van Der Ven-Jongekrijg J, et al. Drosomycin-like defensin, a human homologue of *Drosophila melanogaster* drosomycin with antifungal activity. Antimicrob Agents Chemother. 2008;52: 1407–1412. doi:10.1128/AAC.00155-07

79. Schmitt P, Rosa RD, Destoumieux-Garzón D. An intimate link between antimicrobial peptide sequence diversity and binding to essential components of bacterial membranes. Biochim Biophys Acta - Biomembr. 2016;1858: 958–970. doi:10.1016/j.bbamem.2015.10.011

80. Liu G, Tian Y, Hanson MA, Sah PK, Li J, Lemaitre B. *Drosophila* host defense mechanisms against filamentous fungal pathogens with diverse lifestyles. PLoS Pathog. 2026;22: e1013995. doi:10.1371/journal.ppat.1013995

81. Araki M, Kurihara M, Kinoshita S, Awane R, Sato T, Ohkawa Y, et al. Anti-tumour effects of antimicrobial peptides, components of the innate immune system, against haematopoietic tumours in *Drosophila* mxc mutants. Dis Model Mech. 2019;12. doi:10.1242/dmm.037721

82. Krautz R, Khalili D, Theopold U. Tissue-autonomous immune response regulates stress signaling during hypertrophy. eLife. 2020;9: e64919. doi:10.7554/eLife.64919

83. Swanson LC, Rimkus SA, Ganetzky B, Wassarman DA. Loss of the Aantimicrobial Ppeptide Metchnikowin protects against traumatic brain injury outcomes in *Drosophila melanogaster*. G3: Genes Genomes Genet. 2020;10: 3109–3119. doi:10.1534/g3.120.401377

84. Ferrandon D, Jung AC, Criqui M, Lemaitre B, Uttenweiler-Joseph S, Michaut L, et al. A drosomycin-GFP reporter transgene reveals a local immune response in *Drosophila* that is not dependent on the Toll pathway. EMBO J. 1998;17: 1217–27. doi:10.1093/emboj/17.5.1217

85. Buchon N, Broderick NA, Poidevin M, Pradervand S, Lemaitre B. *Drosophila* intestinal response to bacterial infection: Activation of host defense and stem cell proliferation. Cell Host Microbe. 2009;5: 200–211. doi:10.1016/j.chom.2009.01.003

86. Deng X-J, Yang W-Y, Huang Y-D, Cao Y, Wen S-Y, Xia Q-Y, et al. Gene expression divergence and evolutionary analysis of the drosomycin gene family in *Drosophila melanogaster*. J Biomed Biotechnol. 2009;2009: 315423. doi:10.1155/2009/315423

87. Cohen L, Moran Y, Sharon A, Segal D, Gordon D, Gurevitz M. Drosomycin, an innate immunity peptide of *Drosophila melanogaster*, interacts with the aly voltage-gated sodium channel. J Biol Chem. 2009;284: 23558–23563. doi:10.1074/jbc.M109.023358

88. Amien S, Kliwer I, Márton ML, Debener T, Geiger D, Becker D, et al. Defensin-like ZmES4 mediates pollen tube burst in maize via opening of the potassium channel KZM1. PLoS Biol. 2010;8: e1000388. doi:10.1371/journal.pbio.1000388

89. Okuda S, Tsutsui H, Shiina K, Sprunck S, Takeuchi H, Yui R, et al. Defensin-like polypeptide LUREs are pollen tube attractants secreted from synergid cells. Nature. 2009;458: 357–361. doi:10.1038/nature07882

90. Verster KI, Cinege G, Lipinszki Z, Magyar LB, Kurucz Eâ, Tarnopol RL, et al. Evolution of insect innate immunity through domestication of bacterial toxins. Proc Natl Acad Sci USA. 2023;120: e2218334120. doi:10.1073/pnas.2218334120

91. Tarnopol RL, Tamsil JA, Cinege G, Ha JH, Verster KI, Aâ brahám E, et al. Experimental horizontal transfer of phage-derived genes to *Drosophila* confers innate immunity to parasitoids. Curr Biol. 2025;35: 514–529.e7. doi:10.1016/j.cub.2024.11.071

92. Liu Z, Tao M, Xu Z, Zhang J, Li Y, Dong Z, et al. A bacterial gene acquired by parasitoid wasps contributes to venom secretion against host defence. EMBO J. 2026 [cited 18 Feb 2026]. doi:10.1038/s44318-026-00702-6

93. Brown AN, Lloyd VK. Evidence for horizontal transfer of *Wolbachia* by a *Drosophila* mite. Exp Appl Acarol. 2015;66: 301–311. doi:10.1007/s10493-015-9918-z

94. Rizki RM, Rizki TM. Parasitoid virus-like particles destroy *Drosophila* cellular immunity. Proc Natl Acad Sci USA. 1990;87: 8388–8392. doi:10.1073/pnas.87.21.8388

95. Boudinot BE, Fikáček M, Lieberman ZE, Kusy D, Bocak L, Mckenna DD, et al. Systematic bias and the phylogeny of Coleoptera—A response to Cai et al. (2022) following the responses to Cai et al. (2020). Syst Entomol. 2023;48: 223–232. doi:10.1111/syen.12570

96. Beg A, Baldwin Jr. A. The IkB proteins: multifunctional regulators of Rel/NF-kB transcription factors. Genes Dev. 1993;7: 2064–2070. doi:10.1101/gad.7.11.2064

97. Schreiber J, Jenner RG, Murray HL, Gerber GK, Gifford DK, Young RA. Coordinated binding of NF-κB family members in the response of human cells to lipopolysaccharide. Proc Natl Acad Sci USA. 2006;103: 5899–5904. doi:10.1073/pnas.0510996103

98. Busse MS, Arnold CP, Towb P, Katrivesis J, Wasserman SA. A κB sequence code for pathway-specific innate immune responses. EMBO J. 2007;26: 3826–3835. doi:10.1038/sj.emboj.7601798

99. Collins CF, Alston BT, Hibdige SGS, Raimondeau P, Baker ER, Sotelo G, et al. Regulatory features determine the evolutionary fate of laterally acquired genes in plants. Mascagni F, editor. Mol Biol Evol. 2026;43: msag042. doi:10.1093/molbev/msag042

100. Graham LA, Davies PL. Horizontal gene transfer in vertebrates: A fishy tale. Trends in Genetics. 2021;37: 501–503. doi:10.1016/j.tig.2021.02.006

101. Dong X, Limjunyawong N, Sypek EI, Wang G, Ortines RV, Youn C, et al. Keratinocyte-derived defensins activate neutrophil-specific receptors Mrgpra2a/b to prevent skin dysbiosis and bacterial infection. Immunity. 2022;55: 1645–1662.e7. doi:10.1016/j.immuni.2022.06.021

102. Akoh-Arrey T, Brooks JF. Antimicrobial peptides and proteins as rheostats of intestinal homeostasis and immunity. Curr Opin Immunol. 2026;99: 102738. doi:10.1016/j.coi.2026.102738

103. Arias-Rojas A, Frahm D, Hurwitz R, Brinkmann V, Iatsenko I. Resistance to host antimicrobial peptides mediates resilience of gut commensals during infection and aging in *Drosophila*. Proc Natl Acad Sci USA. 2023;120: e2305649120. doi:10.1073/pnas.2305649120

104. Masson F, Brown RL, Vizueta J, Irvine T, Xiong Z, Romiguier J, et al. Pathogen-specific social immunity is associated with erosion of individual immune function in an ant. Nat Commun. 2024;15: 9260. doi:10.1038/s41467-024-53527-4

105. Login FH, Balmand S, Vallier A, Vincent-Monégat C, Vigneron A, Weiss-Gayet M, et al. Antimicrobial peptides keep insect endosymbionts under control. Science. 2011;334: 362–365. doi:10.1126/science.1209728

106. Brooks JF, Behrendt CL, Ruhn KA, Lee S, Raj P, Takahashi JS, et al. The microbiota coordinates diurnal rhythms in innate immunity with the circadian clock. Cell. 2021;184: 4154–4167.e12. doi:10.1016/j.cell.2021.07.001

107. Nyholm SV, McFall-Ngai MJ. A lasting symbiosis: how the Hawaiian bobtail squid finds and keeps its bioluminescent bacterial partner. Nat Rev Microbiol. 2021;19: 666–679. doi:10.1038/s41579-021-00567-y

108. Khan I, Agashe D, Rolff J. Early-life inflammation, immune response and ageing. Proc R Soc B: Biol Sci. 2017;284: 20170125. doi:10.1098/rspb.2017.0125

109. Candille SI, Kaelin CB, Cattanach BM, Yu B, Thompson DA, Nix MA, et al. A β-defensin mutation causes black coat color in domestic dogs. Science. 2007;318: 1418–1423. doi:10.1126/science.1147880

110. Huang W, Baliga C, Aleksandrova EV, Atkinson G, Polikanov YS, Vázquez-Laslop N, et al. Activity, structure, and diversity of Type II proline-rich antimicrobial peptides from insects. EMBO Rep. 2024. doi:10.1038/s44319-024-00277-5

111. Gagnon MG, Roy RN, Lomakin IB, Florin T, Mankin AS, Steitz TA. Structures of proline-rich peptides bound to the ribosome reveal a common mechanism of protein synthesis inhibition. Nucleic Acids Res. 2016;44: 2439–2450. doi:10.1093/nar/gkw018

112. Hamilton PT, Peng F, Boulanger MJ, Perlman SJ. A ribosome-inactivating protein in a *Drosophila* defensive symbiont. Proc Natl Acad Sci USA. 2016;113: 350–355. doi:10.1073/pnas.1518648113

113. Rozewicki J, Li S, Amada KM, Standley DM, Katoh K. MAFFT-DASH: Integrated protein sequence and structural alignment. Nucleic Acids Res. 2019;47: W5–W10. doi:10.1093/nar/gkz342

114. Sun H, Cole M, Gomes-Ragnoni E, Johnson J, Jarvis A, Hédelin L, et al. Life history and infection susceptibility parameters of *Drosophila* species reared on a common diet. bioRxiv; 2025. p. 2025.11.25.690557. doi:10.1101/2025.11.25.690557

115. Copley RR, Totrov M, Linnell J, Field S, Ragoussis J, Udalova IA. Functional conservation of Rel binding sites in drosophilid genomes. Genome Res. 2007;17: 1327–1335. doi:10.1101/gr.6490707

116. Hanson MA, Cohen LB, Marra A, Iatsenko I, Wasserman SA, Lemaitre B. The *Drosophila* Baramicin polypeptide gene protects against fungal infection. PLoS Pathog. 2021;17: e1009846. doi:10.1371/journal.ppat.1009846

117. Kück P, Meusemann K. FASconCAT: Convenient handling of data matrices. Mol Phylogenetics Evol. 2010;56: 1115–1118. doi:10.1016/j.ympev.2010.04.024

118. Suyama M, Torrents D, Bork P. PAL2NAL: Robust conversion of protein sequence alignments into the corresponding codon alignments. Nucleic Acids Res. 2006;34: W609–W612. doi:10.1093/nar/gkl315

119. Ronquist F, Teslenko M, Mark P, Ayres DL, Darling A, Höhna S, et al. MrBayes 3.2: Efficient bayesian phylogenetic inference and model choice across a large model space. Syst Biol. 2012;61: 539–542. doi:10.1093/sysbio/sys029

120. Morel B, Williams TA, Stamatakis A, Szöllősi GJ. AleRax: A tool for gene and species tree co-estimation and reconciliation under a probabilistic model of gene duplication, transfer, and loss. Bioinformatics. 2024;40: btae162. doi:10.1093/bioinformatics/btae162

121. Morel B, Kozlov AM, Stamatakis A, Szöllősi GJ. GeneRax: A Tool for species-tree-aware maximum likelihood-based gene family tree inference under gene duplication, transfer, and loss. Mol Biol Evol. 2020;37: 2763–2774. doi:10.1093/molbev/msaa141

122. Baril T, Galbraith J, Hayward A. Earl Grey: A fully automated user-friendly transposable element annotation and analysis pipeline. Mol Biol Evol. 2024;41: msae068. doi:10.1093/molbev/msae068

123. Westlake H, Hanson MA, Lemaitre B. The *Drosophila* immunity handbook. First. EPFL Press; 2024. doi:10.55430/6304TDIHVA01

124. Jasso-Martínez JM, Castañeda-Osorio R, Mil-Salazar DA, Pintos-Garduño KF. Phylogeny of Orthoptera (Insecta) using targeted ultraconserved element data. Mol Phylogenetics Evol. 2025;211: 108402. doi:10.1016/j.ympev.2025.108402

125. De Moya RS, Yoshizawa K, Walden KK, Sweet AD, Dietrich CH, Kevin P J. Phylogenomics of parasitic and nonparasitic lice (Insecta: Psocodea): combining sequence data and exploring compositional bias solutions in next generation data sets. Syst Biol. 2021;70: 719–738. doi:10.1093/sysbio/syaa075

126. Peters RS, Krogmann L, Mayer C, Donath A, Gunkel S, Meusemann K, et al. Evolutionary history of the Hymenoptera. Curr Biol. 2017;27: 1013–1018. doi:10.1016/j.cub.2017.01.027

127. Motyka M, Kusy D, Arias ET, Bocak L. Phylogenomics-based click-beetle classification tackles multiple origins of phenotypic modifications. Syst Entomol. 2026;51: e70017. doi:10.1111/syen.70017

128. Zhang H, Song N, Yin X. Higher-level phylogeny of Chrysomelidae based on expanded sampling of mitogenomes. PLOS ONE. 2022;17: e0258587. doi:10.1371/journal.pone.0258587

