## Supplementary material for "Horizontal transfer of an antimicrobial peptide across insects": Data File S: Data File S1.docx

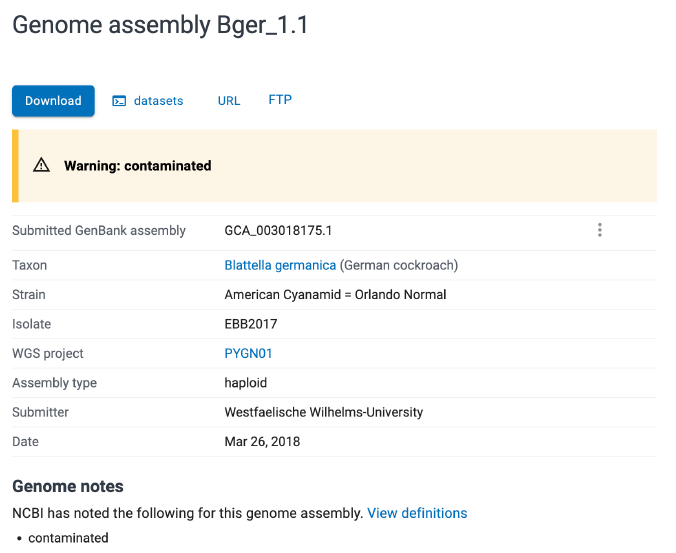

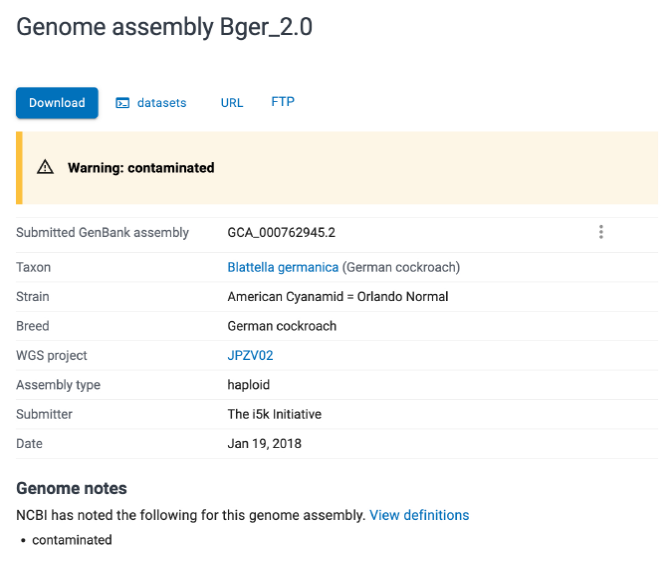

**Data File SX. Screenshot of the NCBI contamination warning of the two *Blattella germanica* genome assemblies**. *Drs* genes were found in the German cockroach *Blattella germanica* in previous studies but available NCBI assemblies of this species are reportedly contaminated (GCA_003018175.1 and GCA_000762945.2). Screenshot taken on 14.05.2025.
