## Supplementary figures and images for "Horizontal transfer of an antimicrobial peptide across insects"

### FigureS1.png

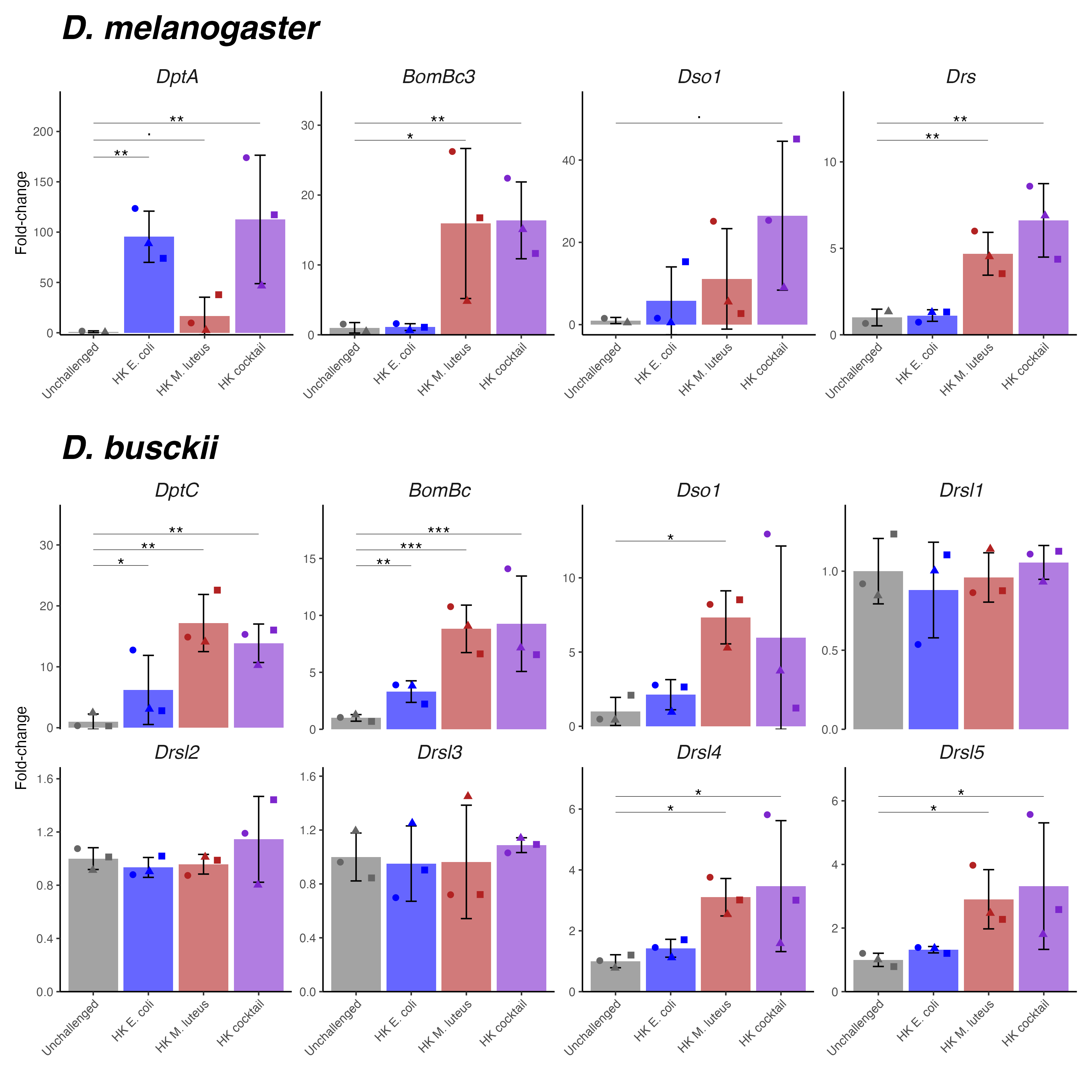

### FigureS2.pdf

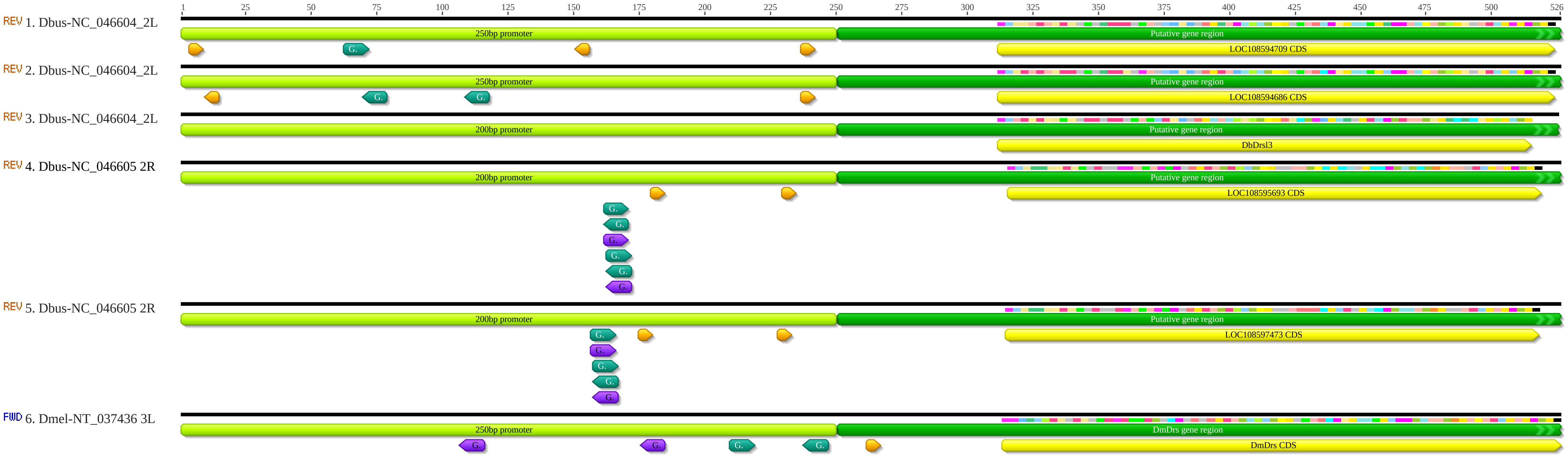

### FigureS3.pdf

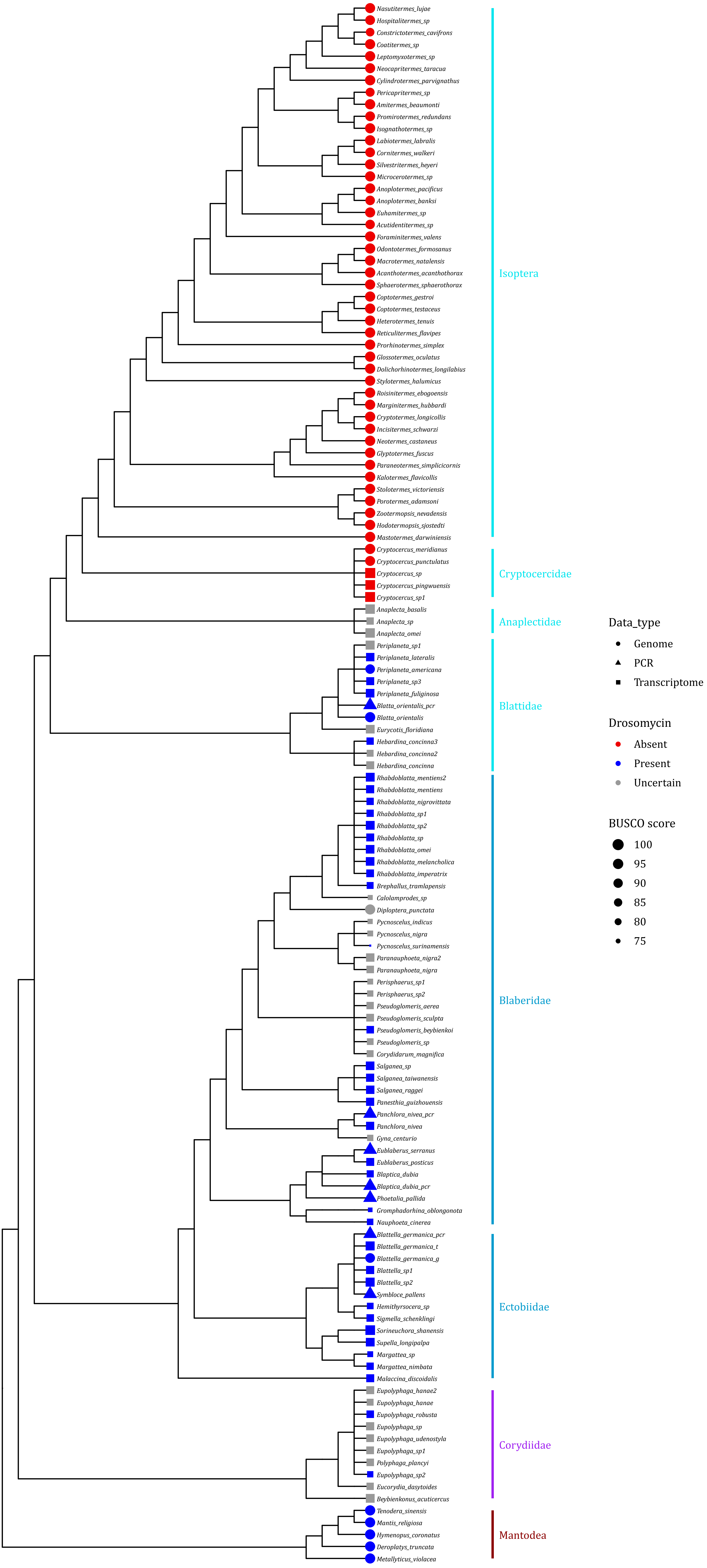

### FigureS4_booklice.pdf

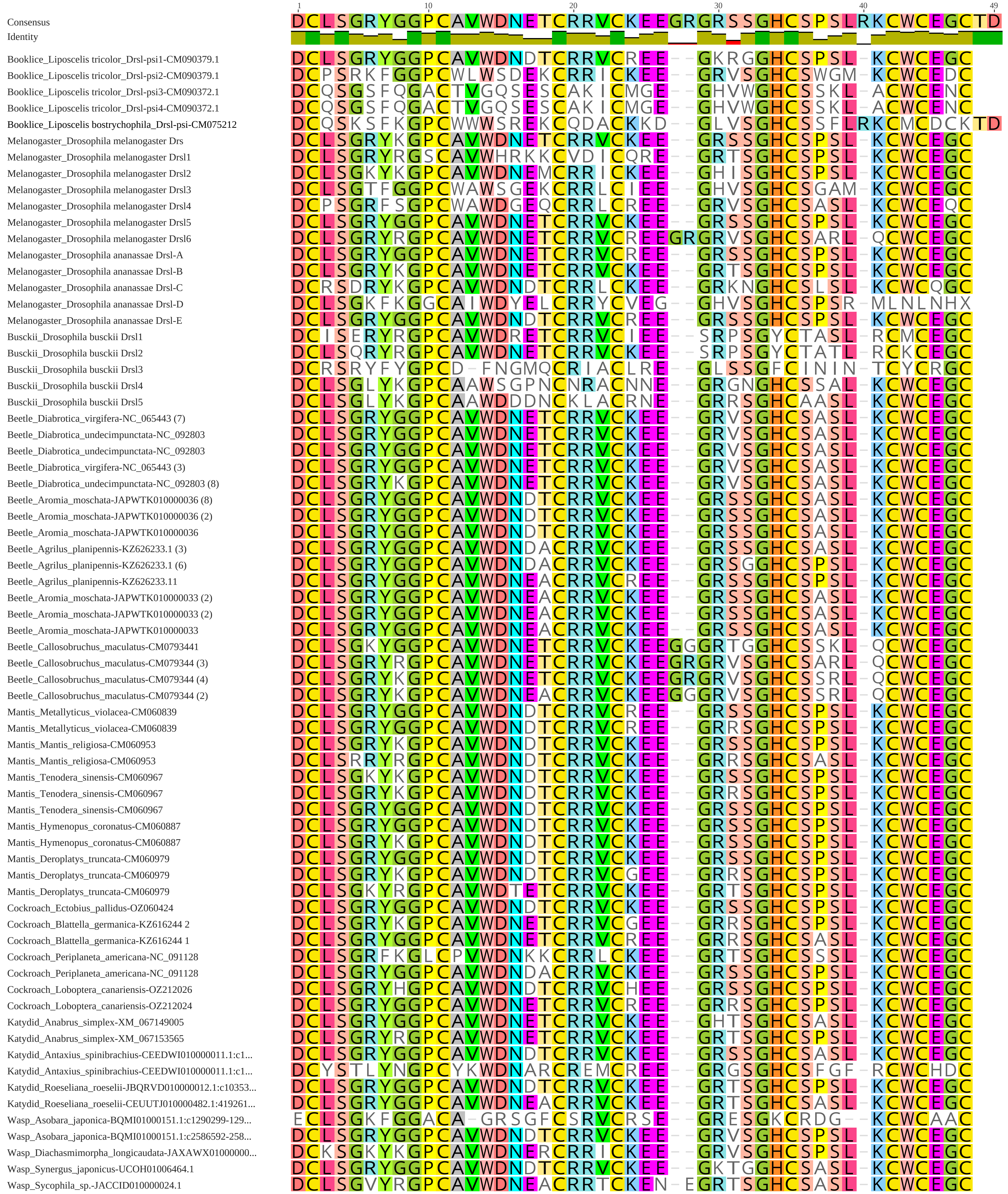

### FigureS5 Full_Drs_Parsimony-20261014.pdf

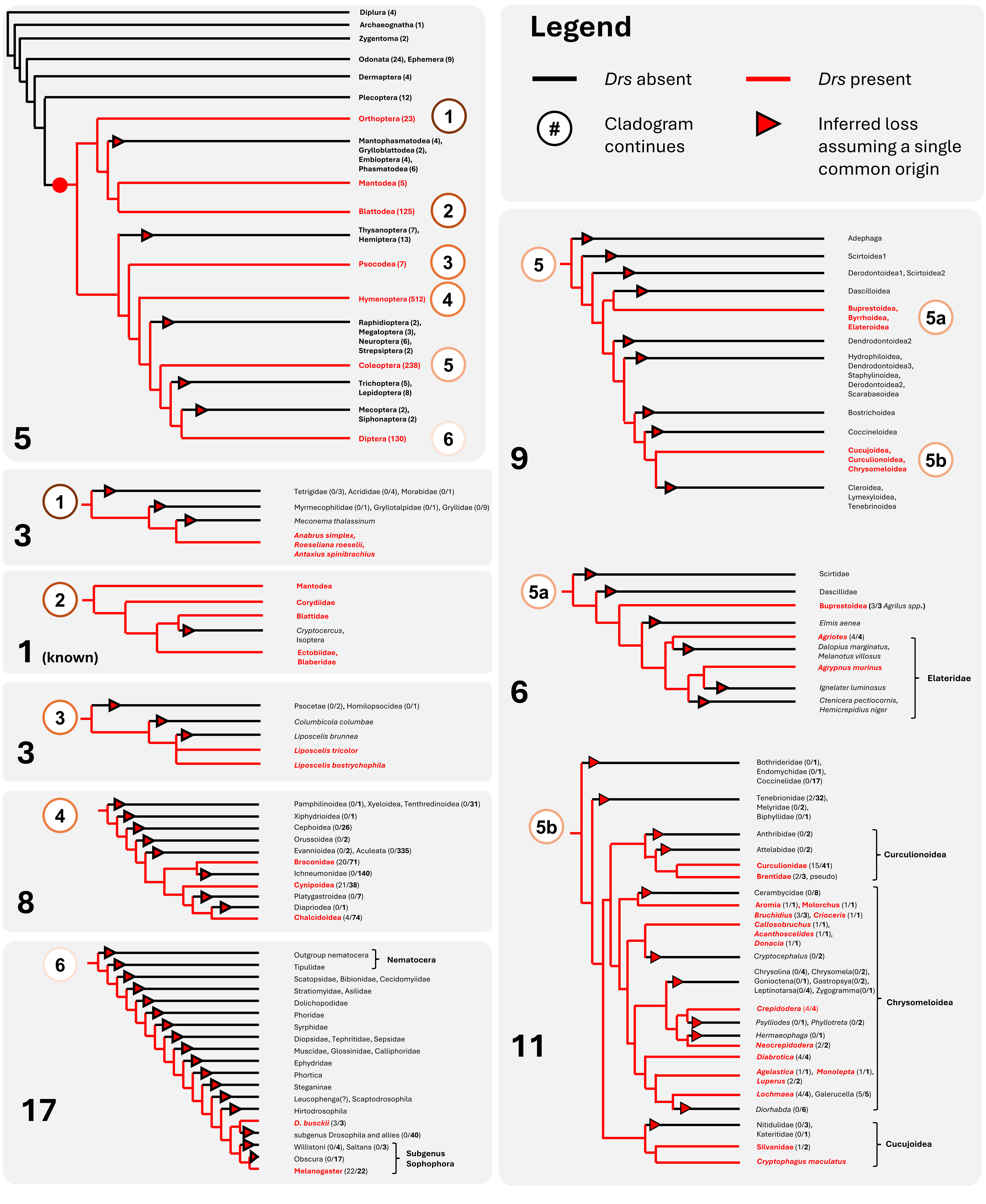

### FigureS6.pdf

a

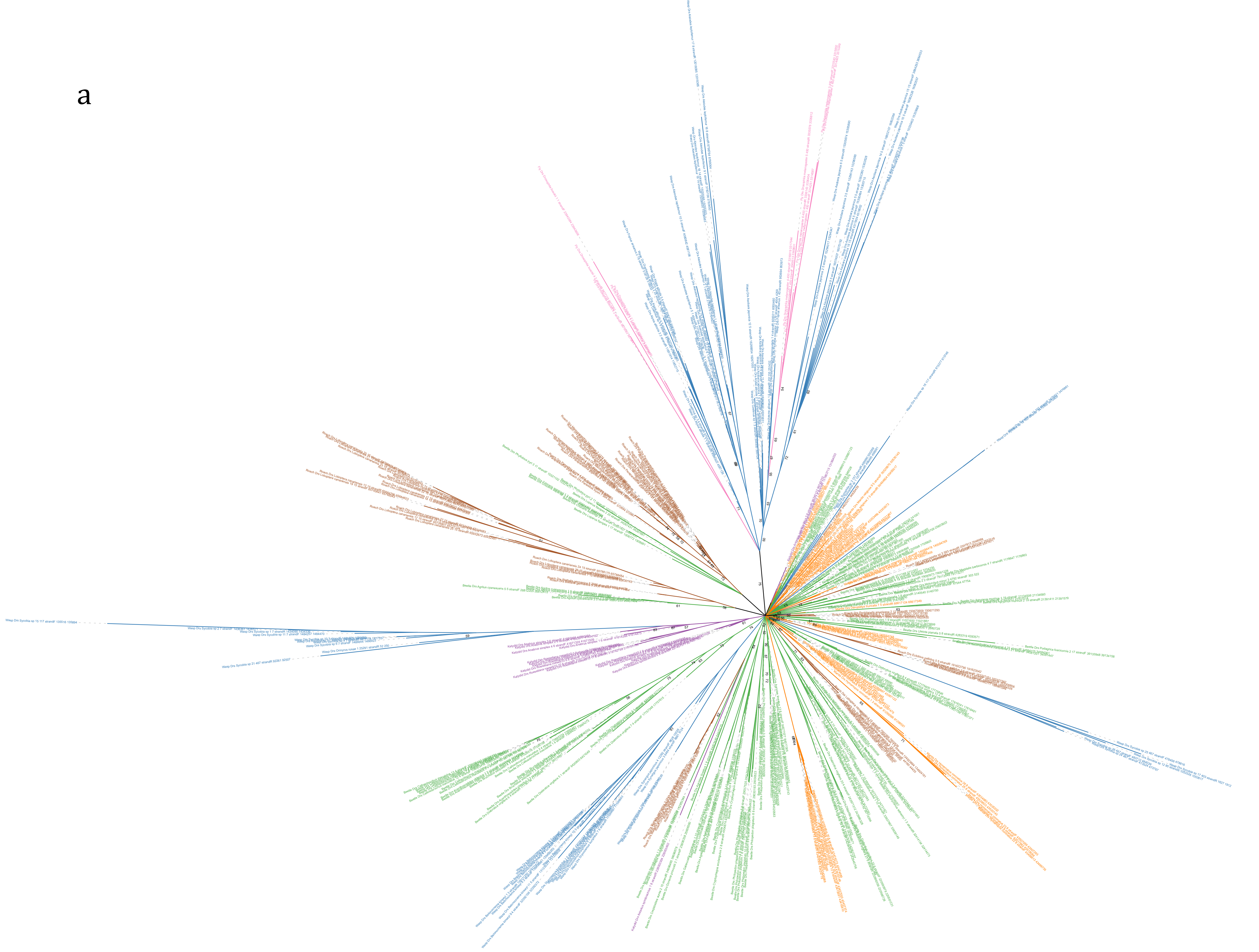

b

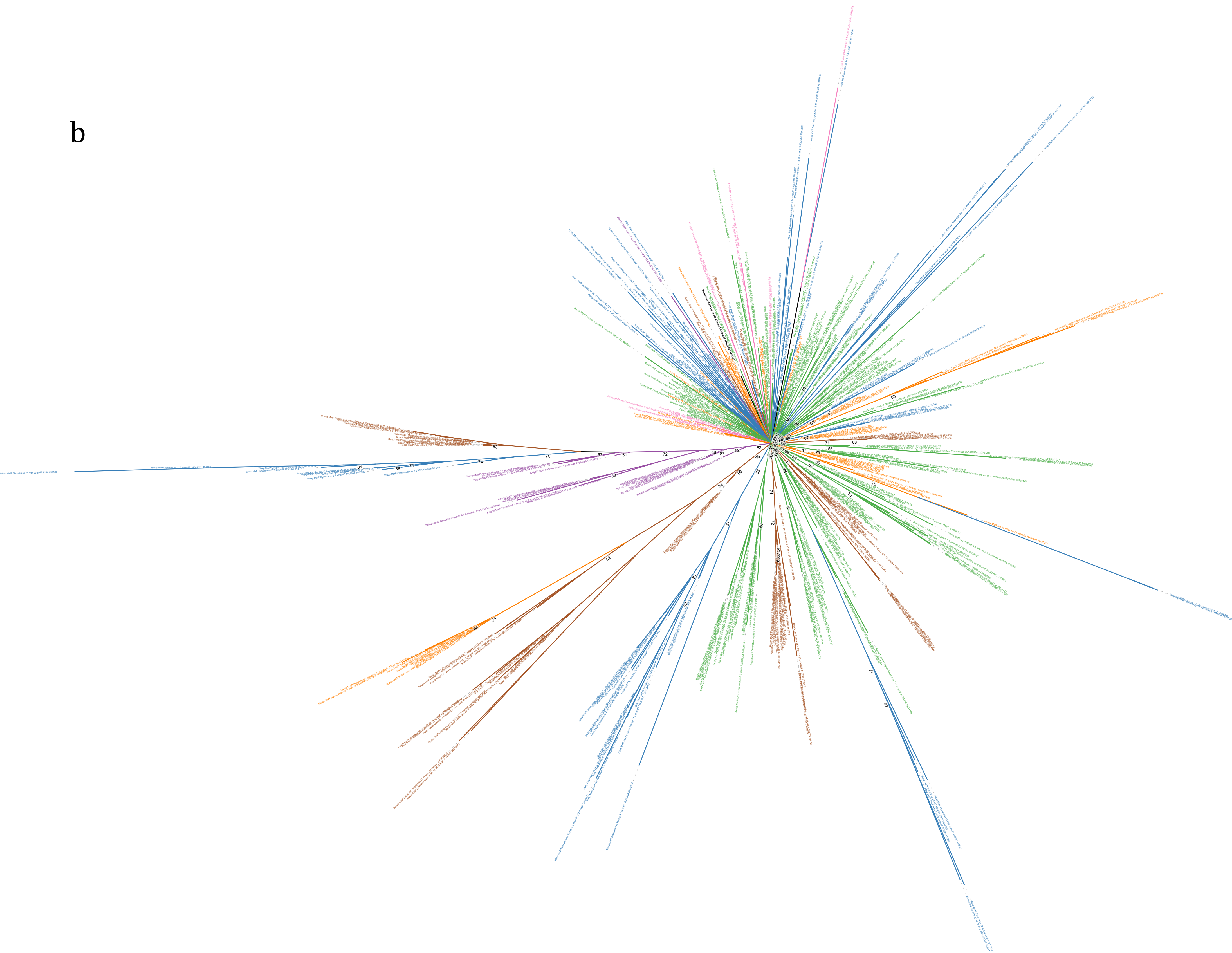

### FigureS7.pdf

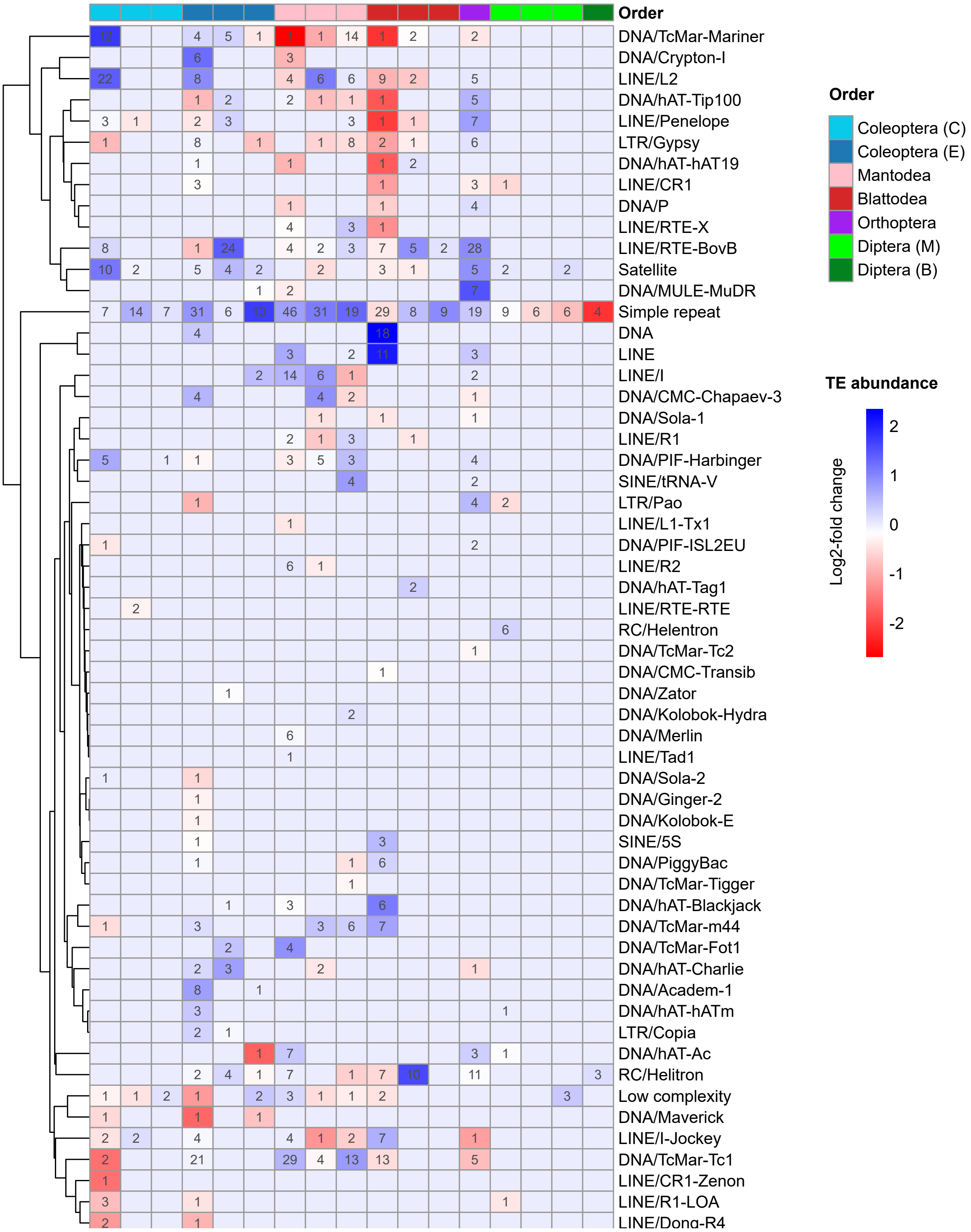

### FigureS8.png

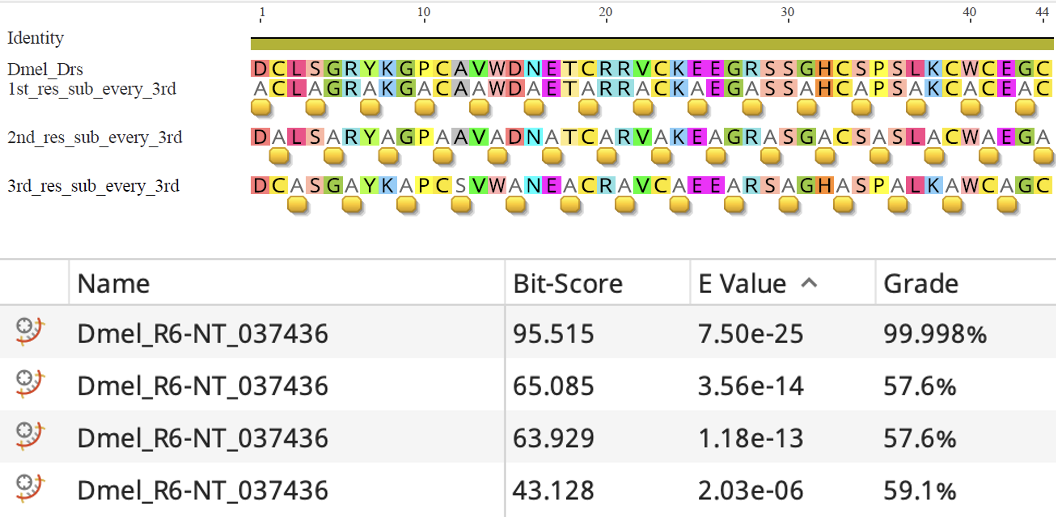
